# Phase separation of YAP reorganizes genome topology for long-term, YAP target gene expression

**DOI:** 10.1101/438416

**Authors:** Danfeng Cai, Daniel Feliciano, Peng Dong, Eduardo Flores, Martin Gruebele, Natalie Porat-Shliom, Shahar Sukenik, Zhe Liu, Jennifer Lippincott-Schwartz

## Abstract

Yes-associated Protein (YAP) is a transcriptional co-activator that regulates cell proliferation and survival by binding to a selective set of enhancers for potent target gene activation, but how YAP coordinates these transcriptional responses is unknown. Here, we demonstrate that YAP forms liquid-like condensates in the nucleus in response to macromolecular crowding. Formed within seconds of hyperosmotic stress, YAP condensates compartmentalized YAP’s DNA binding cofactor TEAD1 along with other YAP-related transcription co-activators, including TAZ, and subsequently induced transcription of YAP-specific proliferation genes. Super-resolution imaging using Assay for Transposase Accessible Chromatin with photoactivated localization microscopy (ATAC-PALM) revealed that YAP nuclear condensates were areas enriched in accessible chromatin domains organized as super-enhancers. Initially devoid of RNA Polymerase II (Pol II), the accessible chromatin domains later acquired Pol II, producing newly transcribed RNA. Removal of YAP’s intrinsically-disordered transcription activation domain (TAD) prevented YAP condensate formation and diminished downstream YAP signaling. Thus, dynamic changes in genome organization and gene activation during YAP reprogramming is mediated by liquid-liquid phase separation.

---

Nuclear localization of the transcriptional co-activator YAP and activation of the TEA domain family (TEAD) transcription factors promote cell proliferation, differentiation and stem cell fate^1–3^, with deficiencies in YAP regulation leading to developmental defects and cancer progression^1^. YAP’s nuclear localization is regulated by the Hippo pathway, an evolutionarily conserved signaling pathway that responds to mechano-chemically induced changes in tissue architecture and osmolarity^4–6^. When the Hippo pathway is active, occurring in densely-plated cells with abundant cell-cell adhesions, YAP binds to 14-3-3 proteins and is retained in the cytoplasm. When the Hippo pathway is inactive, occurring in hyperosmotic conditions or sparsely-plated cells with negligible cell-cell adhesions, YAP is freed from binding to 14-3-3 proteins and redistributes into the nucleus where it triggers transcription of proliferation-specific genes^1,7,8^. Despite YAP’s importance in controlling cell fate and cancer progression, little is known regarding the precise mechanisms that lead to alteration of gene expression patterns when YAP is redistributed into the nucleus.

Phase separation of proteins and nucleic acids into liquid-like or gel-like condensates (also called ‘droplets’) in cells arises through a process involving weak multi-valent interactions of these components^9–11^. A growing body of work supports the role of liquid-liquid phase separation in the formation of distinct sub-nuclear compartments in the nucleus. Phase separation of the protein HP1α, for example, has been shown to contribute to heterochromatin formation^12,13^. Other studies have shown that multiple transcription factors and cofactors can phase separate in the nucleus and up-regulate transcription^14–16^. It has also been proposed that the process of liquid-liquid phase separation leads to clustering of nuclear enhancer elements and bound transcription-related factors to form sites called super-enhancers (SEs)^17^, which transcribe sets of genes important for cell identity or cancer malignancy^17,18^.

ChIP-seq analysis and other work has suggested that YAP interacts with various transcription elongation factors and can act at SE regions in the genome^19,20^. Given this and the above-mentioned suggestion of liquid-liquid phase separation involvement in SE formation, we investigated whether YAP forms nuclear condensates to activate its gene expression program. Using hyperosmotic stress to trigger YAP nuclear redistribution, we saw YAP form nuclear phase-separated, liquid-like condensates within seconds of treatment. The condensates were enriched in TEAD1, a transcription factor that controls YAP target gene expression, as well as the transcription co-factor WW domain containing transcription regulator 1 (TAZ). YAP condensates were initially devoid of RNA Polymerase II (Pol II), but later recruited Pol II for active gene transcription. The timing of Pol II recruitment corresponded to YAP target gene expression in these cells. Employing the super-resolution imaging technique of ATAC PALM to label accessible chromatin regions (i.e., enhancer regions), we demonstrated that YAP condensates under hyperosmotic stress represented highly clustered enhancer regions (i.e., SEs). Together, these results suggest that YAP’s formation of phase-separated condensates in the nucleus in response to hyperosmotic stress helps reorganize the genome for driving long-term, YAP target gene expression.

## RESULTS

### YAP has an intrinsic ability to phase separate into condensates

Proteins that undergo phase separation into condensates usually contain regions of low complexity that are enriched in arginine residues and small polar residues. These low complexity regions (LCRs) promote the weak protein-protein interactions involved in phase separation by charge-charge, π-π, and cation-π interactions^21,22^. Analysis of YAP’s amino acid sequence using algorithms predictive of disordered regions in proteins revealed YAP has an extended C-terminal low complexity region (LCR) encompassing a trans-activating domain (TAD) (Fig. 1a). TADs in other transcription cofactors have been shown to mediate phase separation^15^.

**Fig. 1.**
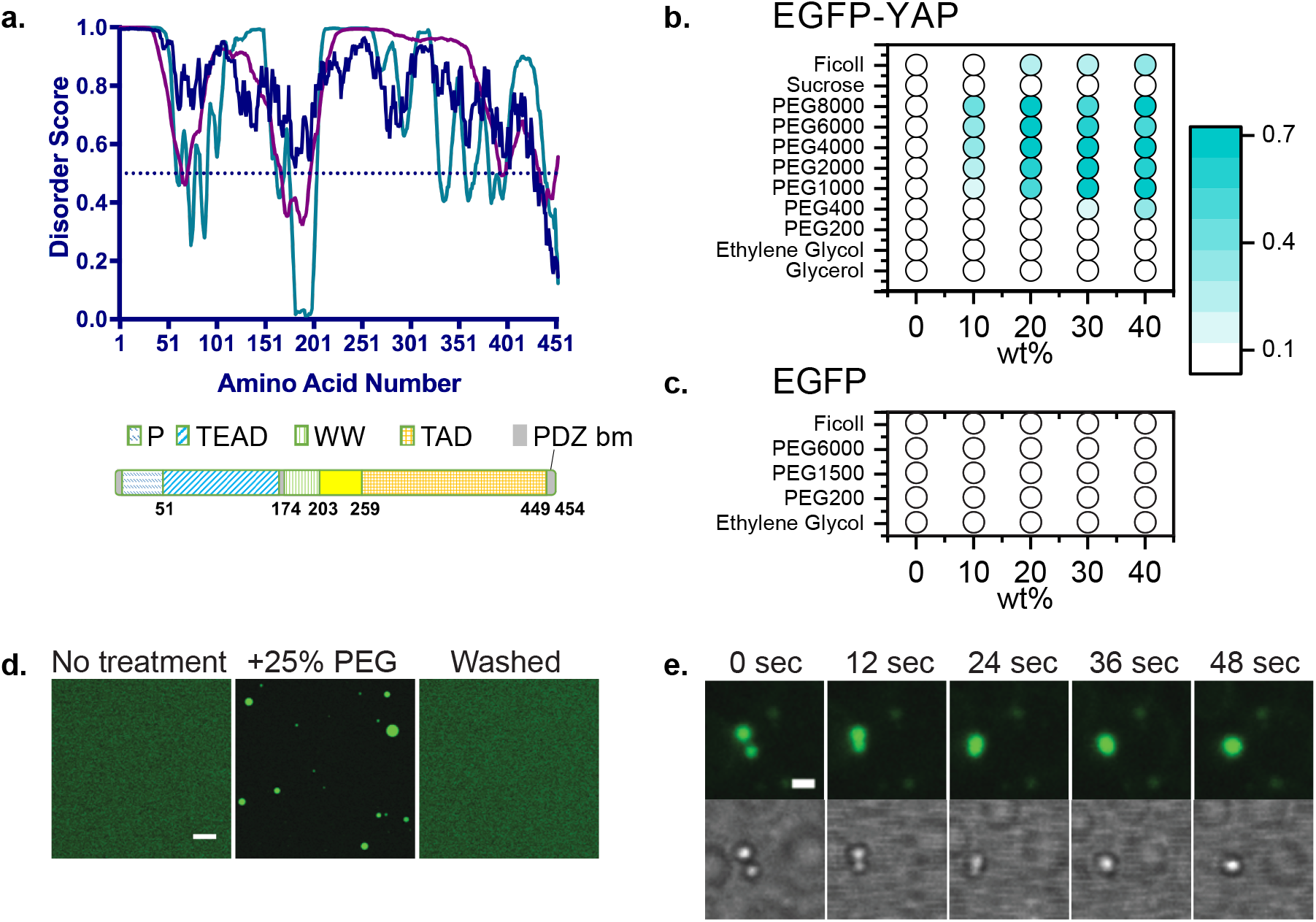
YAP has intrinsic property to phase separate. (a) Disorder analysis of YAP 454aa isoform. Algorithms used: IUPred (blue), VLXT (cyan) and VSL2 (magenta). (b-c) Purified EGFP-YAP shows concentration-dependent increase in turbidity at increasing wt% of large polymeric crowders (b) while purified EGFP does not (c). Color bar: turbidity measured at 600nm (a.u.). (d) EGFP-YAP pre-treatment (left) and after the addition of 25% PEG (middle). Upon centrifugation and resuspension in isotonic buffer, the droplets disappear (right). Scale bar, 10μm. (e) Droplet coalescence shown in fluorescence (top) and brightfield (bottom) illumination. Scale bar, 2μm.

To assess whether YAP has an intrinsic ability to form liquid-like condensates, we purified EGFP-YAP from *E. coli* and examined its biophysical behavior under different conditions. We first measured the phase separation threshold of EGFP-YAP using different solutes (Fig. 1b). In contrast to EGFP protein alone (Fig. 1c), the large, non-ionic crowder such as PEG in sizes larger than 300 Da produced a turbid EGFP-YAP solution (turbidity was also impacted by the concentrations of EGFP-YAP and PEG). Confocal, epifluorescent, and DIC imaging of turbid EGFP-YAP solutions revealed that micron-sized droplet formation occurs in all conditions, and the process was reversible even upon exposure to high PEG concentrations (Fig. 1d). The spheres displayed phase-separated condensate characteristics^23^, including droplet coalescence (Fig. 1e), wetting the coverslip (Movie S1) and the ability to disperse upon resuspension in buffer with no crowder (Fig. 1d, washed). Other conditions known to affect phase separation of proteins (i.e., salts, inert proteins, and nucleic acids)^23,24^ did not trigger YAP droplet formation (Supplementary Fig. 1a). Indeed, varying the salt concentration between 0 and 0.5 M failed to change the turbidity of EGFP-YAP solution, suggesting that strong electrostatic interactions don’t play an important role in YAP phase separation. To see if the intrinsically-disordered TAD domain of YAP is required for YAP phase separation, we constructed an EGFP-YAPΔTAD protein, and found that this mutant underwent phase separation only at concentrations much higher than EGFP-YAP (Supplementary Fig. 1b). These data indicated that YAP has an intrinsic ability to partition into phase-separated condensates through a mechanism that is independent of salts or nucleic acids but dependent on macromolecular crowding, and its TAD region promotes its phase partitioning in vitro.

### YAP undergoes phase separation into cytoplasmic and nuclear condensates in hyper-osmotically stressed cells

YAP is known to redistribute into the nucleus to drive transcription of proliferation-specific genes under conditions such as hyperosmotic stress^6^. Supporting an effect of hyperosmotic stress on YAP target gene transcription, we found that within 3 h of adding the hyperosmotic reagent *D*-sorbitol (0.2M) to HEK293T cells (derived from osmotically-sensitive kidney cells), transcription of YAP target genes Ctgf and Cyr61 increased by 6-fold and 3-fold, respectively (Fig. 2a). Similar results were observed using a different hyperosmotic reagent, PEG300, examined 3 h after treatment (Fig. 2b).

**Fig. 2.**
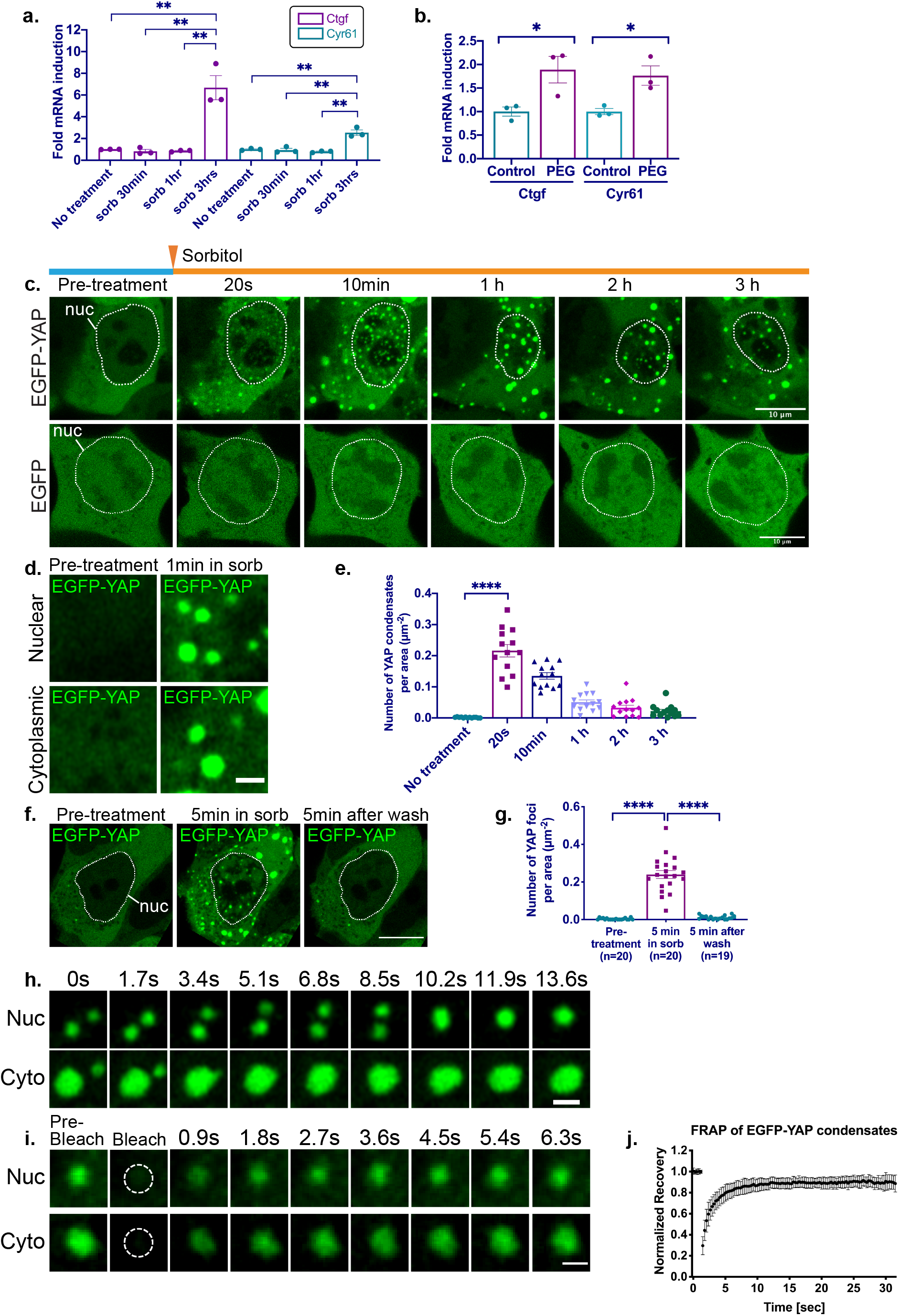
YAP undergoes phase separation under hyperosmotic stress. (a) Quantification of relative Ctgf and Cyr61 mRNAs levels in control and sorbitol-treated HEK293T cells expressing EGFP-YAP. Error bars are SEM. **p<0.01. (b) Quantification of relative Ctgf and Cyr61 mRNAs levels in control and 5% PEG 300-treated HEK293T cells expressing EGFP-YAP. 3hrs after indicated treatments. Error bars are SEM. *p<0.05 (c) EGFP-YAP localizes to the nucleus of HEK293T cells and forms cytoplasmic and nuclear foci 20 sec after sorbitol treatment, while EGFP doesn’t. Scale bars: 10μm. (d) Zoomed-up live-cell imaging of nuclear and cytoplasmic YAP condensates formed after 1 min in sorbitol. Scale bar: μm. (e) Quantification of number of EGFP-YAP foci in the HEK293T cell at indicated time after sorbitol treatment normalized by cell area. Error bars are SEM. (f) Live-cell imaging showing EGFP-YAP condensate formation is reversible. Scale bar: 10μm. (g) Quantification of reversible EGFP-YAP condensate formation. Error bars are SEM. ****p<0.0001. (h) Time-lapse imaging of fusion of nuclear and cytoplasmic EGFP-YAP droplets. (i) FRAP recovery images and quantification of nuclear or cytoplasmic EGFP-YAP condensates. Dotted circle: Spot of photobleaching. Scale bars in (h) and (i) are 1μm. (j) FRAP recovery curve of EGFP-YAP condensates, averaged over 26 experiments. Error bars show SD. Dotted lines indicate the location of the nucleus of HEK293T cells.

Given the above findings, we examined how hyperosmotic stress impacts the distribution of EGFP-YAP in HEK293T cells. Immediately after the addition of *D*-sorbitol, EGFP-YAP underwent a dramatic redistribution from being uniformly dispersed in the cytoplasm to being concentrated in discrete puncta in the cytoplasm and nucleus (Fig. 2c, top panel). The puncta formed within 20s of sorbitol treatment, and in zoomed-in images appeared spherical (Fig. 2d). This contrasted with the behavior of EGFP expressed in these cells (Fig. 2c, lower panel), which showed no change in subcellular distribution upon hyperosmotic stress. The EGFP-YAP puncta persisted for 1 h and then decreased in number until a few nuclear condensates were left after 3 h in sorbitol (Fig. 2c, upper panel; Fig. 2e). Washing in isotonic medium when the puncta were still present caused the puncta to completely disappear within 5 min (Figs. 2f, g).

The EGFP-YAP puncta in hyperosmotically stressed cells exhibited key features of phase-separated condensates^9,22^. This included having high sphericity in high resolution images (Supplementary Fig. 2a), the ability to fuse with each other to form larger spheres in both the nucleus and the cytoplasm (Fig. 2h, Movie S2-S4), and exhibiting rapid fluorescence recovery upon photobleaching (t_1/2_ = 0.9 ± 0.4 sec, Figs. 2i, j). These results, combined with our above finding that the puncta rapidly disassembled upon return to isotonic medium, suggested that the YAP foci formed under hyperosmotic stress represented phase-separated, liquid-like condensates rather than solid aggregates.

The formation of YAP condensates was YAP isoform-independent (Supplementary Fig. 2b) and not limited to the specific osmotic agent used (Supplementary Fig. 2c). Furthermore, examining EGFP-YAP behavior in other cells that were not kidney-derived, including HeLa and mouse embryonic fibroblast (MEF) cells, revealed YAP condensates also formed in these cells in response to hyperosmotic stress (Supplementary Fig. 2d). Thus, condensate induction was independent of cell type. As expected, all of the cells when exposed to hyperosmotic stress exhibited volume reduction in both the nucleus and the cytoplasm (Supplementary Fig. 2e) and increased macromolecular crowding (Supplementary Figs. 2f, g).

### Endogenous YAP in cells forms condensates

To address whether EGFP-YAP resembles the behavior of endogenous YAP protein, we used an antibody to label endogenous YAP in HEK293T cells. Within 10 min of sorbitol treatment, endogenous YAP was localized to the nucleus, appearing in small puncta (Fig. 3a). Like EGFP-YAP, these condensates dissipated after 3 h of treatment. Knocking down YAP by RNAi reduced YAP protein expression measured by immunoblotting and immunofluorescence (Supplementary Figs. 3a, b) and eliminated endogenous YAP foci formation in response to sorbitol treatment (Fig. 3b). Endogenous YAP thus showed similar behavior as EGFP-YAP in HEK293T cells.

**Fig. 3.**
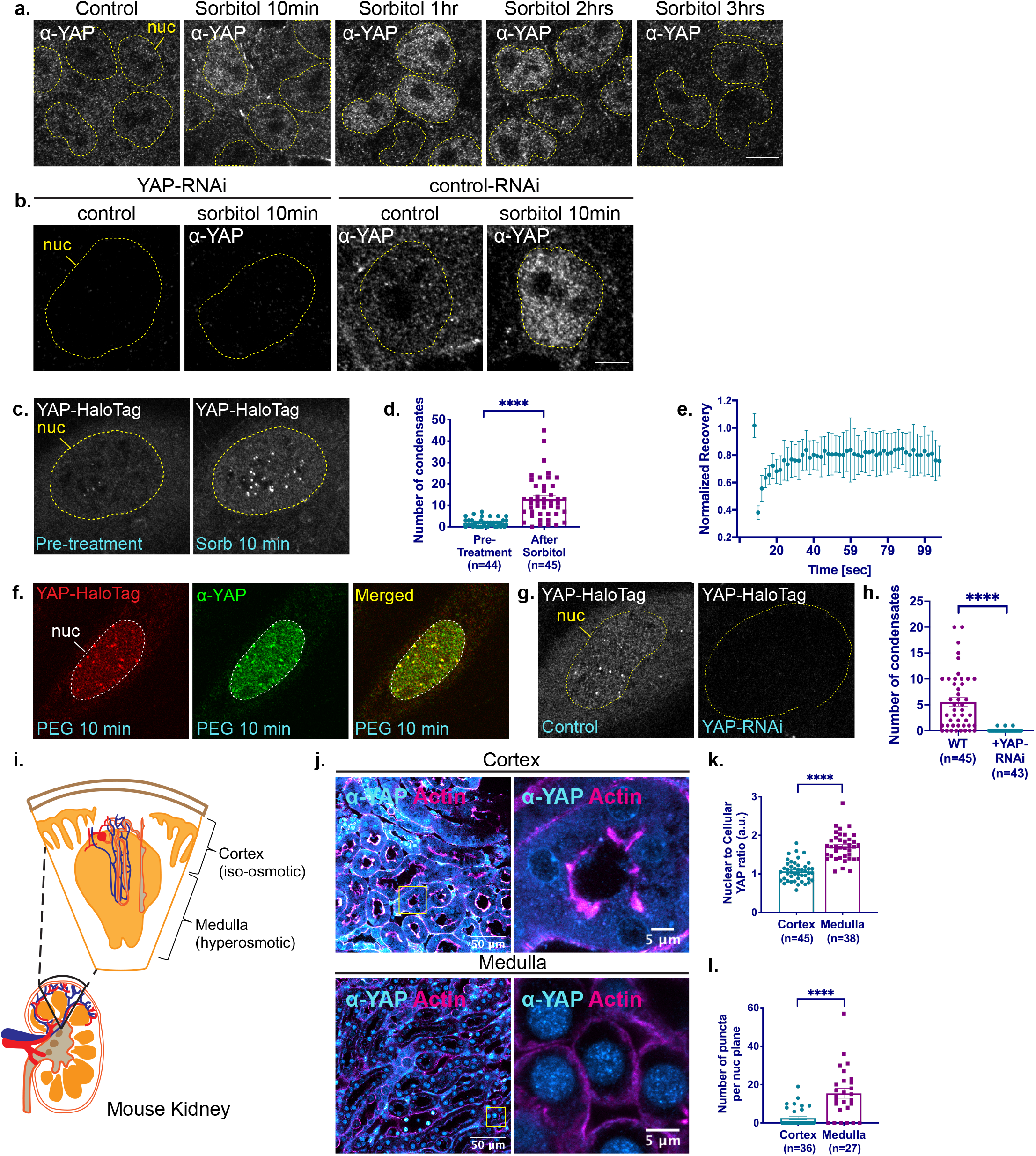
Endogenous YAP form condensates. (a) Endogenous YAP in HEK293T cells visualized by immunofluorescence using antibody staining, showing rapid localization of endogenous YAP to the nucleus. (b) Immunofluorescence images showing endogenous YAP in HEK293T cells form condensates in response to 0.2M sorbitol treatment for 10 min. YAP signal in both diffuse and condensate form is depleted by siRNA against YAP. (c-h) Endogenous Halo-tagged YAP form condensates in U-2 OS cells, visualized by Airyscan live-cell imaging. Live-cell imaging (c) and quantification (d) showing nuclear Halo-YAP condensate labelled by JF549 Halo dye increases in number after sorbitol treatment. (e) FRAP recovery curve of YAP-HaloTag condensate in sorbitol-treated cells. (f) Immunofluorescence showing YAP-HaloTag (labelled with JF549 Halo dye) colocalizes with endogenous YAP labeled with YAP antibody in cells treated with 10% PEG300 for 10 min. (g-h) Live-cell imaging by Airyscan (g) and quantifications (h) showing YAP RNAi knocks down YAP-HaloTag signal in both diffuse and condensate form. Scale bars are 10μm. Error bars show SEM. Unpaired t-test analysis. *p<0.05, **p<0.01, ****p<0.0001. (i-l) Medulla region cells in the mouse kidney have nuclear YAP in condensate form. (i) Illustration of a kidney and relative locations of cortex and medulla. (j) Representative YAP immunofluorescence images of mouse kidney cells at cortex and medulla regions. Larger field of view shown on left, and the yellow boxed regions are magnified and shown on right. (k) Quantification of nuclear to cellular ratio of YAP in indicated regions. (l) Quantification of number of condensates in kidney cells with visible condensates. ***p<0.001, ****p<0.0001.

To characterize endogenous YAP dynamics in another cell line, we generated a CRISPR knock-in U-2 OS cell line, fusing HaloTag to the C-terminus of the YAP genomic loci to generate cells with endogenous YAP replaced with YAP-HaloTag (Supplementary Fig. 3c). YAP-HaloTag nuclear puncta appeared in these cells within 10 min of hyperosmotic stress induced by either 0.2 M sorbitol (Figs. 3c, d) or 10% PEG300 (Supplementary Figs. 3d, e). The puncta recovered rapidly after photobleaching (Fig. 3e) and could be decorated with anti-YAP antibodies (Fig. 3f), suggesting they were phase-separated condensates rather than protein aggregates or artefacts of CRISPR labeling. As expected, knocking down YAP with siRNA to remove YAP-HaloTag/YAP within these cells, resulted in no appearance of anti-YAP labeled condensates (Figs. 3g, h).

We next used immunofluorescence labeling to examine YAP localization in cells from the cortex and medulla region of a mouse kidney, sites that experience iso-osmotic and hyperosmotic conditions, respectively^25–28^ (Fig. 3i). YAP antibody localization in kidney cells in the cortical region, which experiences less change in osmolarity^25–28^, was restricted primarily to the cytoplasm (Fig. 3j, top; Fig. 3k), with few if any nuclear puncta seen (Figs. 3j, top; Fig. 3l). By contrast, YAP localization in the medulla region, which experiences hyperosmotic conditions, was enriched in the nuclei of these cells (Fig. 3j, bottom; Fig. 3k) and appeared in prominent punctate structures that resembled YAP nuclear condensates seen in osmotically-stressed HEK293T and U-2 OS cells (Figs. 3j, bottom; Fig. 3l). These results suggested that endogenous YAP in a tissue can also form liquid-like condensates when experiencing hyperosmotic stress.

### Characterizing cytoplasmic YAP condensates induced by hyperosmotic stress

Phase-separated condensates provide cells with an additional way to organize their internal space, functioning as hubs to facilitate protein-protein interactions or to sequester proteins away from their normal partners^9^. With this in mind, we examined what other proteins co-segregate with YAP in the YAP-enriched condensates seen under hyperosmotic stress. We focused first on YAP condensates that appeared in the cytoplasm at the onset of hyperosmotic stress. We reasoned that these might be involved in facilitating YAP entry into the nucleus through segregation of kinases modifying YAP for nuclear import. Consistent with this possibility, YAP cytoplasmic condensates were found to concentrate proteins involved in YAP-specific post-translational modifications, including Hippo pathway kinase large tumor suppressor 1 (LATS1) (Fig. 4a) and Nemo-like kinase (NLK) (Fig. 4b). Since prior work has shown that hyperosmotic stress leads to activation of both NLK and LATS1/26, these findings raise the possibility that localization of NLK and LATS1/2 within cytoplasmic YAP condensates might coordinate the activity of both kinases.

**Fig. 4.**
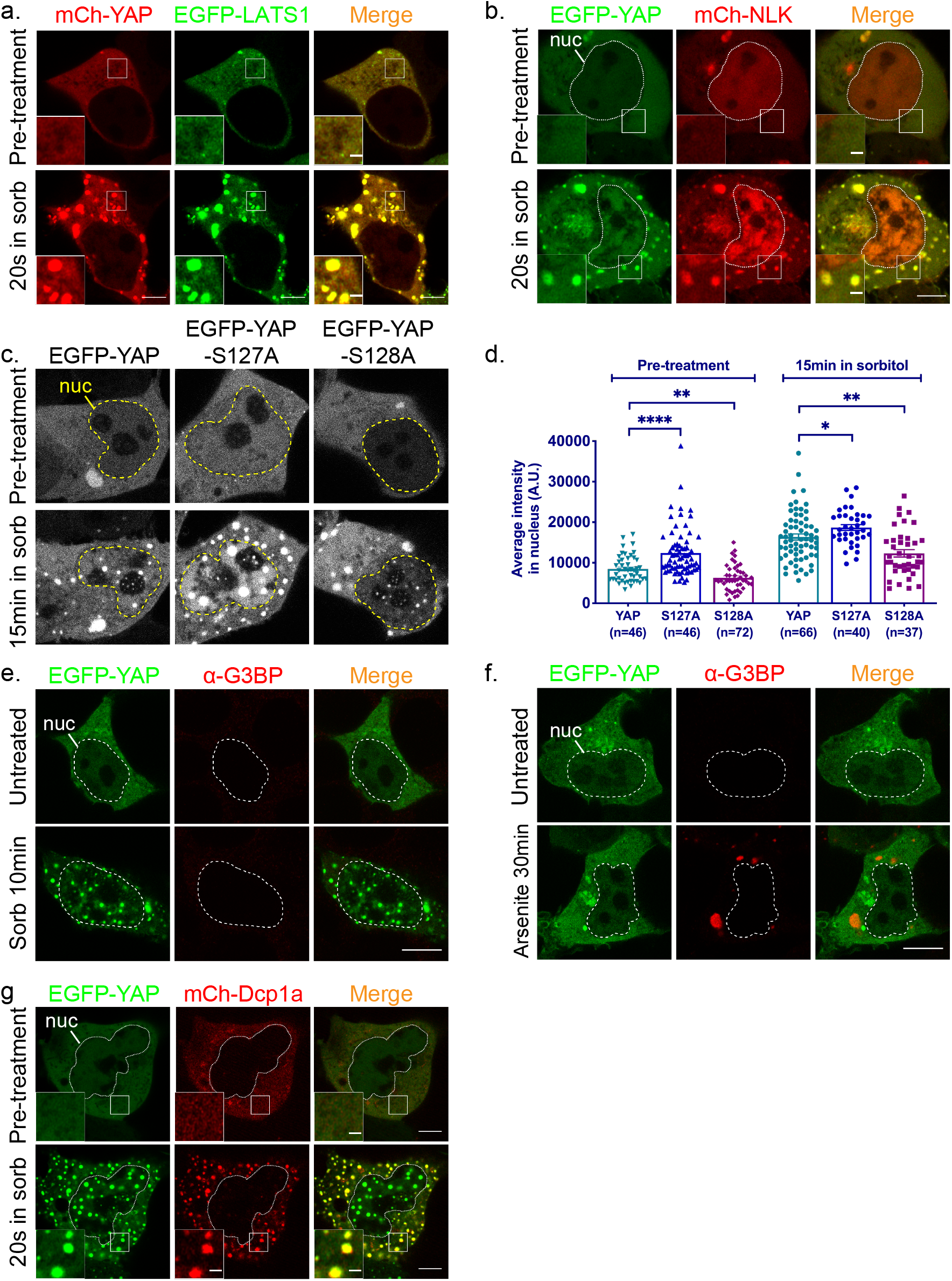
Cytoplasmic EGFP-YAP selectively enrich different proteins. (a) Live-cell imaging showing colocalization of mCherry-YAP condensates with EGFP-LATS1 condensates in the cytoplasm of HEK293T cells after hyperosmotic stress. (b) Live-cell imaging showing colocalization of EGFP-YAP condensates with mCherry-NLK condensates in the cytoplasm of HEK293T cells after hyperosmotic stress. (c) Live-cell imaging of HEK293T cells showing EGFP-YAP wildtype and mutants are able to form condensates with sorbitol treatment, although nuclear localization before and after sorbitol treatment changes, quantified in (d). Comparisons are done between mutants and WT YAP, using unpaired t-test. *p<0.05, **p<0.01, ****p<0.0001. (e) Representative immunofluorescence images showing 0.2M sorbitol treatment for 20 sec induces EGFP-YAP condensate formation in HEK293T cells but not stress granule formation (visualized by G3BP antibody). (f) Representative immunofluorescence images showing 0.5mM arsenite treatment for 30 min induces stress granule formation but not YAP condensate formation. (g) Live-cell imaging showing colocalization of EGFP-YAP condensates with mCherry-Dcp1a condensates in the cytoplasm of HEK293T cells after hyperosmotic stress. In a, b, g: scale bars are 5μm in whole-cell images, and 1μm in magnified views of boxed regions. In e-f: scale bars are 10μm.

LATS1/2 is known to phosphorylate YAP at Ser127, causing it to become more tightly associated with 14-3-3 proteins and retained in the cytoplasm^7,29^. Because there was less mCherry-YAP localized in the nucleus in EGFP-LATS1 overexpressing cells compared to cells not expressing EGFP-LATS1 after sorbitol treatment (compare Fig. 4a and Fig. 2c), the data suggested that enhanced phosphorylation of YAP at Ser127 helps retain YAP in the cytoplasm in EGFP-LATS1 overexpressing cells. Consistent with this, mutating serine 127 to alanine in YAP (to prevent phosphorylation by LATS) significantly increased YAP’s nuclear localization during hyperosmotic shock (Fig. 4d).

NLK phosphorylates YAP at Ser128, releasing YAP from binding to 14-3-3 proteins to allow YAP nuclear import^6,30^. To test whether phosphorylation of YAP at Ser128 in cytoplasmic droplets helps promote YAP’s nuclear redistribution in NLK-expressing cells, we mutated serine 128 to alanine in YAP. We found this significantly impaired YAP’s ability to become nuclear-localized in response to hyperosmotic shock (Fig. 4d), without impairing its ability to form droplets (Fig. 4c).

YAP cytoplasmic droplets did not contain the stress granule component G3BP (Fig. 4e) nor did they co-segregate with G3BP-containing stress granules during arsenite treatment (Fig. 4f). They also were not processing bodies (P-bodies) involved in RNA processing as they lacked critical P-body components, including GW182 and Ago2 (Supplementary Figs. 4a-d). However, the P-body component Dcp1a did co-segregate with YAP droplets (Figs. 4g), suggesting some unknown link to RNA processing. Altogether, the data revealed that YAP cytoplasmic condensates are neither stress granules nor P-bodies, but are a novel type of cytoplasmic droplet that sequesters kinases that target YAP for either cytoplasmic retention or nuclear translocation.

### Characterizing nuclear YAP condensates induced by hyperosmotic stress

We next investigated what molecules co-segregate with YAP condensates in the nucleus, hoping to gain clues as to the roles of YAP in this environment. YAP nuclear condensates did not co-localize with known nuclear body markers such as PML (promyelocytic leukemia bodies, Figs. 5a, b) or Coilin (Cajal bodies, Figs. 5c, d), suggesting they perform functions different from these well-known nuclear bodies. Notably, YAP condensates were enriched in the transcription factor TEAD1 (Figs. 5e, f), which regulates transcription of YAP target genes^1,31,32^. Neither YAP nor TEAD1 were found in nuclear or cytoplasmic droplets prior to sorbitol treatment. Within 2 min of hypertonic stress, however, both molecules were found co-localized in nuclear condensates. Line scans through the condensates showed a similar profile of enrichment for each protein in the droplet (Fig. 5f). A similar co-localization of YAP and TEAD1 in droplets was seen in U-2 OS cells in response to hyperosmotic stress induced by PEG (Supplementary Figs. 5a, b).

**Fig. 5.**
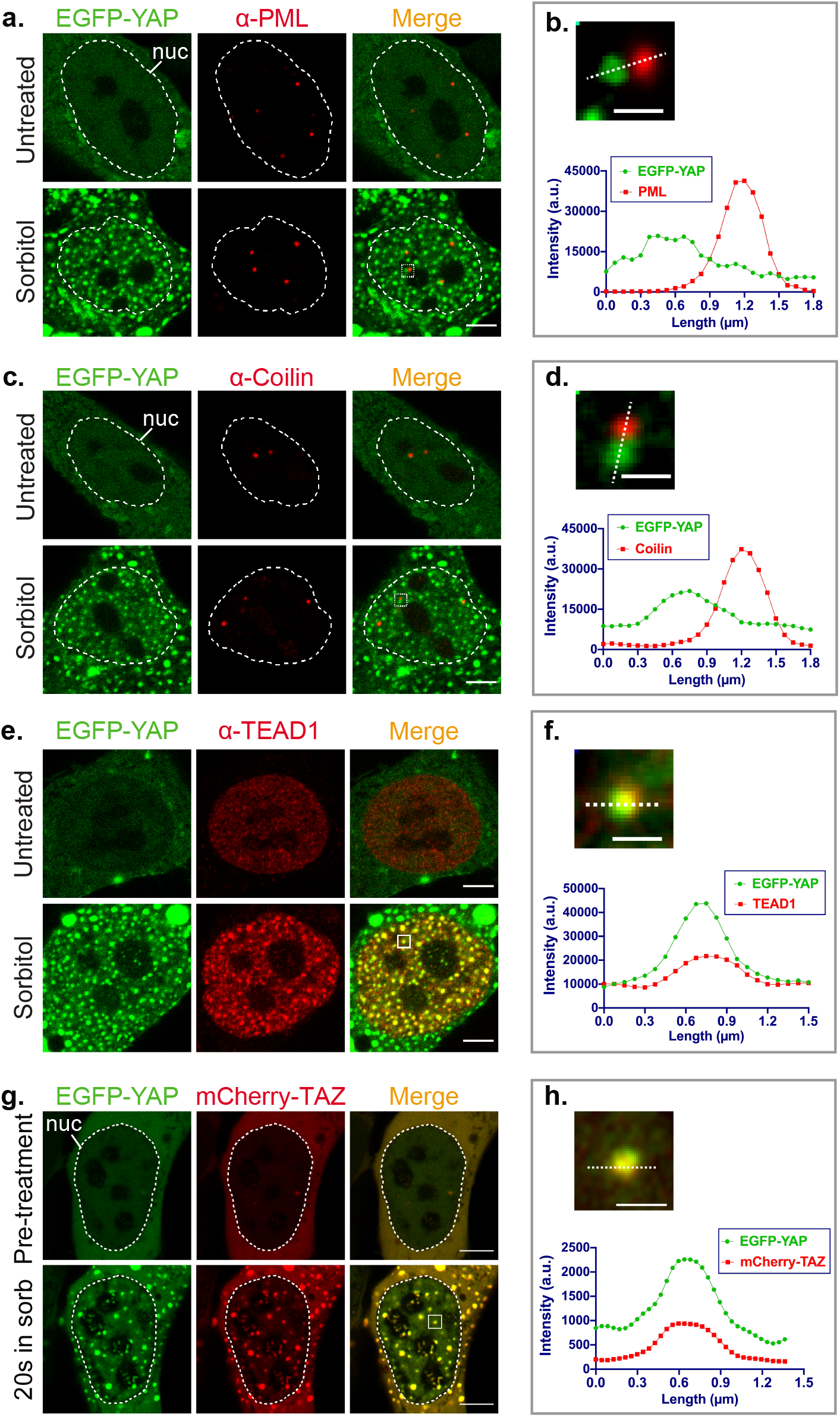
Nuclear EGFP-YAP selectively enrich transcription-related proteins. (a) Immunofluorescence showing no colocalization of PML staining with EGFP-YAP before or after hyperosmotic shock. (b) Magnification of inset of bottom right image of (a) and line-scan. (c-d) Cajal bodies (visualized by anti-Coilin immunofluorescence) don’t colocalize with EGFP-YAP before or after hyperosmotic shock in HEK293T cells. Similar to (a-b). (e) Immunofluorescence showing TEAD1 colocalizes with nuclear EGFP-YAP condensates under hyperosmotic stress. (f) Magnification of the boxed region in (g), and line plot of the dotted line. (g) Live-cell imaging showing colocalization of mCherry-TAZ with EGFP-YAP 20 sec after hyperosmotic stress. (h) Magnification of the boxed region in (j), and line plot of the dotted line. Scale bars are 5μm in a, c, e, g and 1μm in b, d, f, j.

YAP nuclear condensates formed under hyperosmotic stress also contained the paralog of YAP, WW domain containing transcription regulator (TAZ) (Figs. 5g, h). In HEK293T cells, TAZ and YAP both were homogenously dispersed in the cytoplasm prior to hyperosmotic treatment, but redistributed into nuclear droplets within 20 sec of treatment (Fig. 5g). In the droplets, YAP and TAZ showed similar overall distributions (Fig. 5h). As recent work has reported that overexpressed TAZ tagged with a fluorescent protein localizes to nuclear droplets without any hyperosmotic stress^33^, we asked whether YAP redistributes into these droplets upon hypertonic stress. To address this, we co-expressed EGP-TAZ and mCherry-YAP in U-2 OS cells, in which a small pool of EGFP-TAZ, but not mCherry-YAP was seen in nuclear puncta prior to hyperosmotic stress. After hyperosmotic stress, both EGFP-TAZ and mCherry-YAP now co-localized in nuclear condensates, which had significantly increased in number (Supplementary Fig. 5c). Observing the same cell before and after hyperosmotic treatment revealed mCherry-YAP redistributed into pre-existing EGFP-TAZ droplets, as well as into new ones, which EGFP-TAZ then also distributed into (Supplementary Fig. 5c).

### 3D ATAC PALM reveals YAP nuclear condensates localize to spatially-segregated accessible chromatin domains

The linear distribution of chromatin accessible for binding by transcription factors, promoters and insulators has been mapped by DNase I digestion^34,35^ and by assay for transposase-accessible chromatin with high-throughput sequences (ATAC-seq)^36^, revealing that many gene enhancer elements cluster with their transcription factors into super-enhancer (SE) regions. Given that nuclear YAP condensates were enriched in YAP-specific transcription factors and coactivators, we asked whether these condensates marked SE regions. To test this, we visualized accessible chromatin in cells using a recently developed ATAC-based imaging method called 3D ATAC photoactivated localization microscopy (ATAC-PALM)^37^. In this approach, Tn5 transposase is used to insert photo-activatable fluorescent DNA probes into the accessible chromatin domains (Fig. 6a, left), and then photoactivated localization microscopy (PALM)^38^ combined with Lattice Light Sheet microscopy (LLSM)^39,40^ is employed to visualize accessible chromatin regions in 3D through the entire nucleus (Fig. 6a, right).

**Fig. 6.**
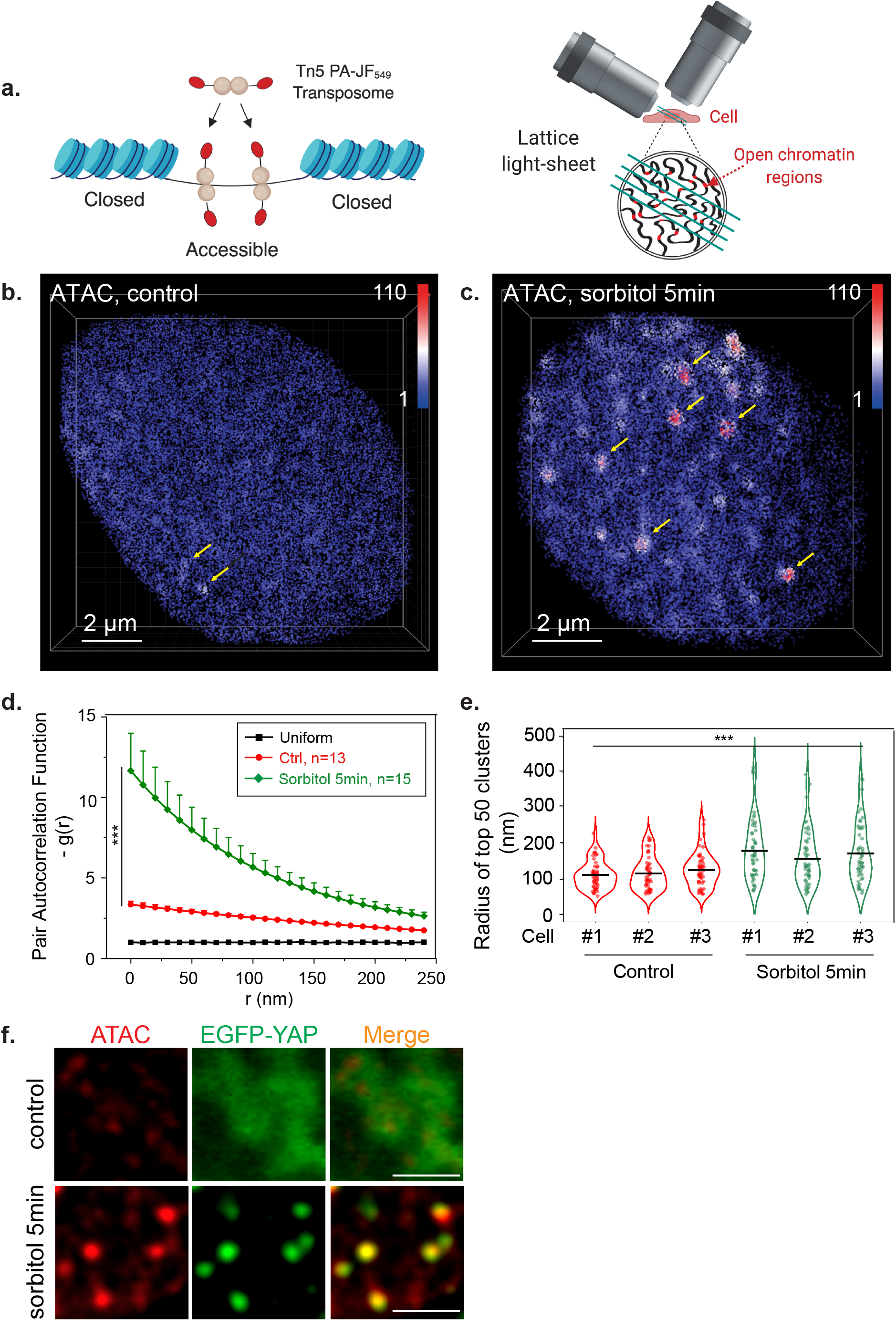
YAP condensates induced by sorbitol treatment alter 3D organization of accessible chromatin. (a) Schematics of the 3D ATAC-PALM microscopy labeling and imaging strategy. Photoactivatable Janelia Fluor 549 (PA-JF549) was conjugated to DNA oligo containing the mosaic ends (ME) of the Tn5 transposon and reconstituted with Tn5 transposase (cyan) to form the active transposome complex in vitro (left panel). The cells were then fixed, permeabilized, and accessible sites in the genome were selectively labeled by the reconstituted active transposome complex before mounting onto the Lattice Light Sheet microscope for 3D ATAC-PALM imaging. (b-c) 3D image of accessible chromatin localizations in control (b) and 5min sorbitol-treated (c) HEK293T cells. The color-coded localization density was calculated with a canopy radius of 250nm. Yellow arrows point to ATAC clusters identified. (d) Global pair autocorrelation function g(r) analysis of ATAC-PALM localizations for control and sorbitol-treated HEK 293 cells. The error bars represent standard error (SE) of the mean. The non-parametric Mann-Whitney U test was used to compare the clustering amplitude A (equals to g(0)) in different groups. g(r) was plotted from the fitted exponential decay function. (e) Normalized radii of identified accessible chromatin clusters in control and sorbitol-treated HEK293T cells identified by DBSCAN algorithm. For statistical test, data from 3 individual cells for each condition were pooled together and Mann-Whitney U test was applied. (f) Example images show colocalization of identified accessible chromatin clusters and YAP condensates in control and sorbitol-treated HEK293T cells. To derive the ATAC-PALM intensity map (red) in order to compare with EGFP-YAP signal, ATAC-PALM localizations were binned within a cubic of 100 nm with a 3D Gaussian filter and a convolution kernel of 3×3×3. Scale bar: 2 μm.

Prior to hyperosmotic shock, ATAC-PALM imaging revealed that accessible-chromatin regions were widely dispersed throughout the nucleus with a few of these regions clustered into observable puncta (Fig. 6b, arrows, Movie S5). Strikingly, after 5 min of hyperosmotic treatment with sorbitol, large ATAC-labeled clusters were seen distributed across the nucleus (Fig. 6c, arrows, Movie S6). Pair autocorrelation analysis, used to quantify differences in the density and size of the clustered accessible-chromatin regions, revealed that accessible-chromatin regions under sorbitol treatment, on average, became significantly more clustered than before treatment (Fig. 6d). Measurement of the radius of the ATAC clusters in sorbitol-treated cells demonstrated that some clusters were over 400 nm in diameter, significantly larger than those in control cells, whose average diameter was ~120 nm (Fig. 6e). The significant size increase in ATAC-labeled clusters could reflect fusion of neighboring accessible chromatin regions, the appearance of entirely new accessible-chromatin regions, or both.

To test whether the enlarged ATAC-positive structures under sorbitol treatment represented YAP condensates, we compared the localizations of ATAC structures with EGFP-YAP condensates in sorbitol-treated cells. Remarkably, virtually all ATAC-positive structures were associated with the EGFP-YAP condensates (Fig. 6f, Movie S7). This suggested that osmotically-driven phase separation of YAP into condensates reorganizes the genome into clusters of accessible chromatin regions enriched in YAP and its binding partners.

### Nuclear YAP condensates become sites of active gene transcription

To determine whether nuclear YAP condensates with accessible chromatin domains are sites of active gene transcription, we probed them with antibodies to RNA Polymerase II (Pol II). RNA Pol II was initially segregated from YAP nuclear droplets at 10 min of hyperosmotic shock using sorbitol (Figs. 7a, b; Supplementary Fig. 7a). However, after 2 h in sorbitol RNA Pol II was no longer excluded from YAP condensates. Instead, RNA Pol II now decorated the surface of nuclear YAP condensates (Fig. 7c, Supplementary Fig. 7b). This suggested that YAP nuclear condensates, while initially driving clustering of gene enhancer elements to form SEs, do not recruit RNA Pol II until later, presumably after the cell has had time to adapt to hyperosmotic conditions for reorganization of its transcription program.

**Fig. 7.**
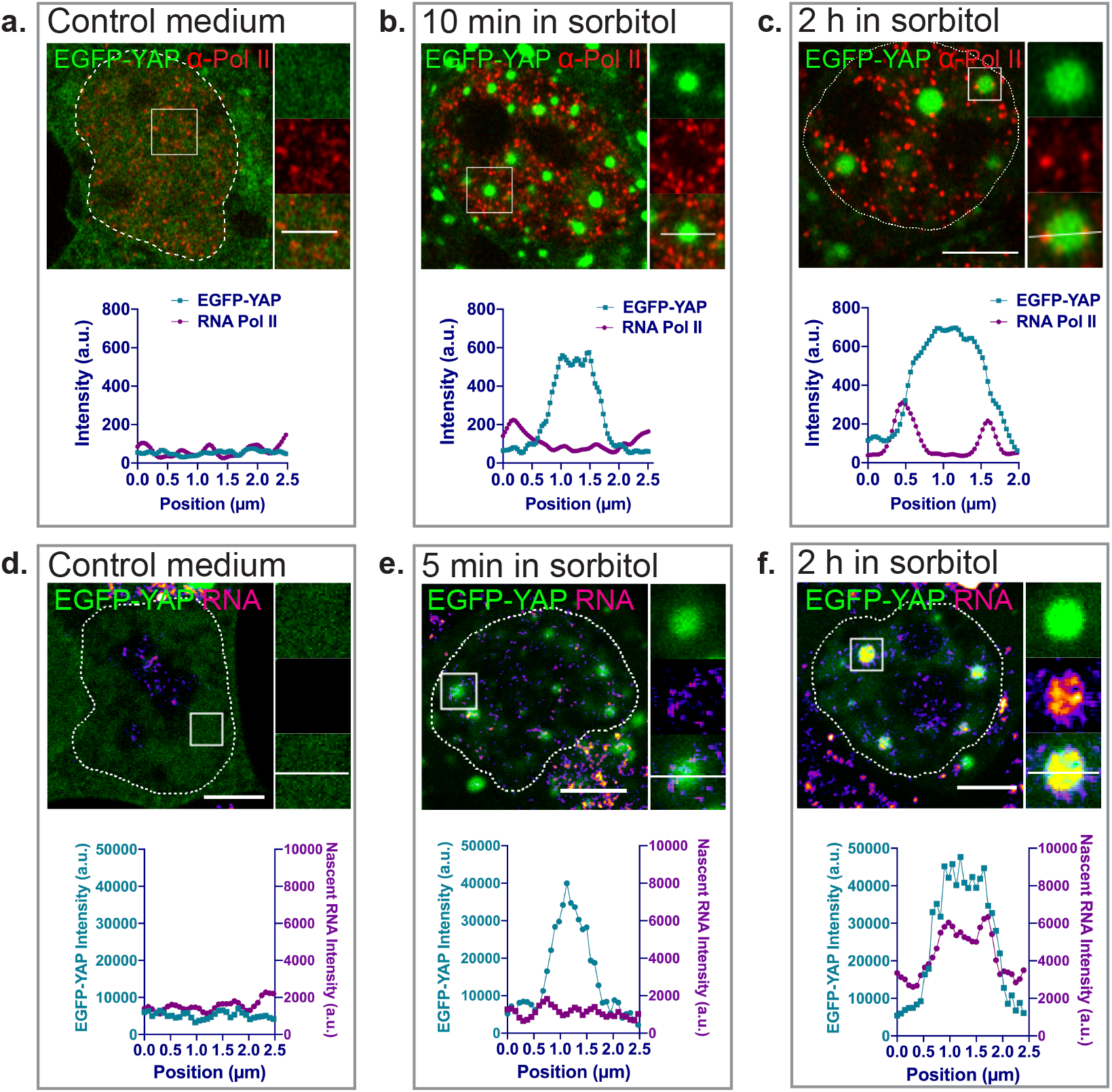
EGFP-YAP nuclear condensate localization with RNA Pol II during sorbitol treatment. (a-c) Immunofluorescence images and line plot of indicated regions showing EGFP-YAP localization with RNA Pol II (pSer2) in different conditions. (d-f) EGFP-YAP localization with nascent RNA pulse-labelled by EU for 5min, and visualized by Click-iT chemistry. RNA is shown in the Fire look-up table in ImageJ. Line plots of EGFP-YAP condensate localization with nascent RNA signal are shown below. Scale bars: 5μm.

Given that RNA Pol II localizes to the surface of YAP condensates by 2 h of hyperosmotic stress, we examined the distribution of nascent RNA to determine whether the condensate sites were active for gene transcription. Nascent RNA did not specifically localize at clustered sites before treatment or at YAP-containing nuclear condensates after 5 min sorbitol treatment (Figs. 7d, e). However, 2 h after cells were in sorbitol, a large amount of nascent RNA was transcribed selectively at clustered sites associated with nuclear YAP condensates (Fig. 7f). That this leads to an increase of YAP target gene transcription was suggested by RT-PCR experiments, which showed an increase in expression of the YAP target gene products Ctgf and Cyr61 within 3 h of sorbitol treatment (Fig. 2a).

### YAP’s intrinsically-disordered transcription activation domain is responsible for YAP condensate formation and its downstream effects on transcription

We next sought to identify and remove regions in YAP responsible for its phase separation under hyperosmotic stress to test whether YAP droplet formation is necessary for the above effects on gene transcription. To identify regions responsible for YAP phase separation, we deleted the intrinsically-disordered C-terminal transcriptional activation domain (TAD) of YAP and then generated an EGFP-YAPΔTAD fusion protein (Fig. 8a). Upon expression of EGFP-YAPΔTAD in cells, we found no EGFP-YAPΔTAD-containing condensates formed upon hyperosmotic shock, in contrast to cells expressing EGFP-YAP (Fig. 8b, c). This suggested that without its TAD sequence YAP is unable to phase separate into condensates. We additionally found that EGFP-YAPΔTAD expression disrupted endogenous YAP’s ability to form condensates in cells (Supplementary Fig. 8b-c). In addition, less EGFP-YAPΔTAD redistributed into the nucleus at 10 min after sorbitol treatment compared to EGFP-YAP (Fig. 8 d). This suggested that expression of YAPΔTAD can interfere with endogenous YAP’s ability both to transition into phase-separated condensates and to become nuclear localized under hyperosmotic stress.

**Fig. 8.**
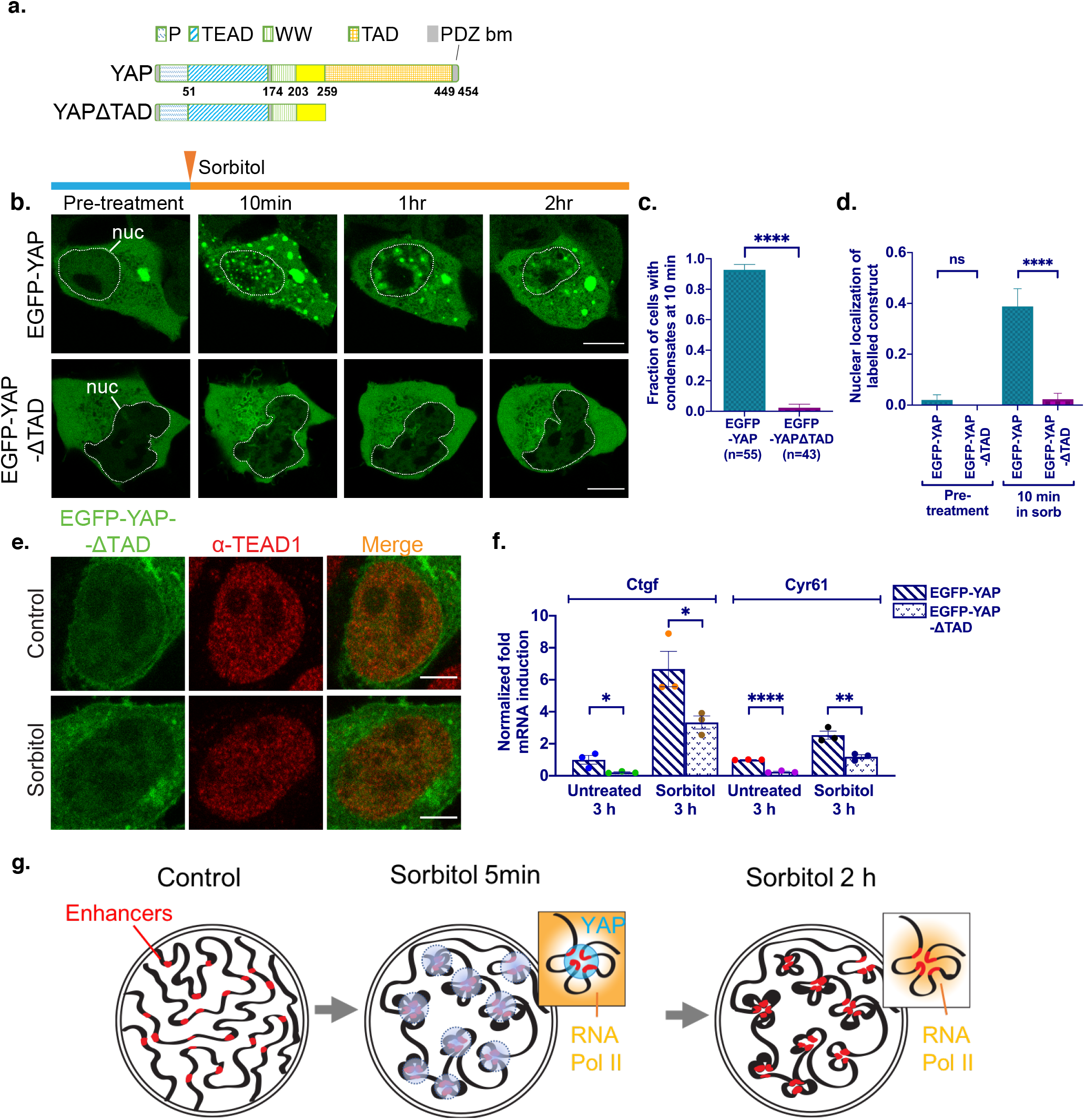
Phase separation of YAP is important for its nuclear localization and signaling. (a) Illustration of wildtype YAP (isoform 3, uniprot identifier: P46937-3) and YAPΓTAD structures. Numbers show amino acid position in the wildtype YAP. (b) Livecell imaging showing time-dependent phase separation of EGFP-YAP variants in HEK293T cells. Scale bars: 10μm. (c) Quantification of fraction of HEK293T cells expressing EGFP-YAP variants with liquid droplets at 10 min after sorbitol treatment. Error bars show SEM. ****p<0.0001. (d) Quantification showing fraction of cells that have nuclear localized EGFP-YAP or EGFP-YAPΓTAD. Nuclear localization means higher or equal average intensity in the nucleus versus the cytoplasm. Error bars show SEM. NS: not significant. ****p<0.0001. (e) Immunofluorescence image of relative localization of EGFP-YAPΓtAd with TEAD1 in control or 0.2M sorbitol-treated HEK293T cells. Scale bars: 5μm. (f) Quantification of RT-PCR results showing relative Ctgf and Cyr61 mRNA expressions in HEK293T cells expressing EGFP-YAP variants after indicated treatments. Error bars show SEM. *p<0.05, **p<0.01, ****p<0.0001. (g) Model depicting remodeling of enhancer elements by YAP phase separation.

We next examined whether YAP phase separation into droplets is necessary for long-term promotion of YAP transcriptional activity. In EGFP-YAPΔTAD expressing cells, in which YAP condensates didn’t form, TEAD1 did not enrich in any nuclear foci upon hyperosmotic stress (Fig. 8e). Measuring Ctgf and Cyr61 mRNA expression 3 h after sorbitol treatment revealed a significant decrease in expression of the mRNAs in cells expressing EGFP-YAPΔTAD compared to cells expressing EGFP-YAP (Fig. 8f). These results suggested that the formation of YAP condensates, and later expression of YAP target genes, during sorbitol treatment is dependent on YAP’s C-terminal TAD domain.

## Discussion

Our work shows that the nuclear localization and activity of YAP, a key Hippo pathway transducer, is regulated by liquid-liquid phase separation. Specifically, hyperosmotic stress-induced condensation of YAP into liquid droplets shifts YAP’s distribution from cytoplasm to nucleus, where it drives transcription of genes involved in cell proliferation. Because cytoplasmic and nuclear YAP droplets sequestered different sets of proteins, with cytoplasmic droplets containing NLK and nuclear droplets containing TEAD1, their formation appeared to contribute both to YAP’s ability to redistribute into the nucleus and to its role in transcriptional control. Our findings thus highlight a new mechanism involving phase separation that enables YAP to collaborate with different factors in different cellular compartments to control its activity during signaling.

The Hippo signaling pathway governs tissue growth and homeostasis by regulating cell proliferation and differentiation, with control of these cellular processes occurring through pathway-mediated localization of the downstream effectors YAP/TAZ. The overall activity of Hippo signaling is to prevent the transcriptional activity of downstream effectors of YAP and TAZ, accomplished by retention of these transcription co-factors in the cytoplasm through the activity of a cascade of core kinases (including the kinases MST1/2 and Lats1/2). While many signals activate the Hippo pathway (i.e., by engagement of receptors that activate the core kinases), the events that antagonize the pathway have only begun to be understood. One antagonizing event is phosphorylation of YAP at Ser 128 by NLK, which occurs in response to hyperosmotic stress and leads to YAP redistribution into the nucleus^6^. Our finding that NLK co-localizes with YAP in cytoplasmic droplets under hyperosmotic stress suggests this co-compartmentalization serves to rapidly alter YAP phosphorylation kinetics through increased kinase- and substrate-concentrations in the droplet. The additional presence of Lats1 in YAP cytoplasmic droplets could serve to modulate this effect, as we observed decreased pools of YAP in the nucleus in Lats1 overexpressing cells. YAP sequestration into cytoplasmic condensates under hyperosmotic stress could help prevent YAP from interacting with 14-3-3 proteins, such as 14-3-3σ protein, which we found did not partition into cytoplasmic YAP droplets (data not shown). In addition, cytoplasmic YAP condensates could also protect YAP from targeted degradation by phosphorylated casein kinase in the cytoplasm^41^. Understanding how these and other core Hippo kinase members, including MST1/2, PP2A, Sav1 and MOB1, distribute and behave when YAP droplets form under hyperosmotic stress is an important area for future work as their modulation could be relevant in therapeutic studies.

Another area impacted by our findings relates to nuclear reorganization by YAP condensates. A growing body of work suggests the importance of phase separation in re-organizing the genome for transcription^14–17^. Tools for studying these processes in live cells, such as the light-induced phase separation systems developed by Brangwynne and colleagues^42–44^, have only recently become available. Our finding that osmotically-driven assembly and later disassembly of YAP nuclear condensates provides a novel, physiologically relevant approach for analyzing condensate-induced changes in nuclear organization. Indeed, using the super-resolution imaging method of ATAC-PALM to visualize accessible chromatin regions, we found that the cell’s genome reorganized dramatically within 5 min of hyperosmotic stress, with dispersed accessible chromatin domains observed in untreated cells becoming highly concentrated in large, YAP-containing nuclear condensates after treatment.

Canonically, YAP binds to TEAD family members to induce transcription of YAP target genes. But how the targeted genomic loci are coordinately controlled has been unclear. We found TEAD1, TAZ and accessible chromatin regions co-localize with newly formed YAP droplets within minutes of hyperosmotic stress. This raises the possibility that YAP nuclear condensates physically pull in targeted genomic loci while pushing out non-targeted regions of neighboring genome to allow for synchronized transcription of YAP target genes, a characteristic of nuclear condensates proposed by others^14,15,17,43^. During this genome reorganization, we observed that while Pol II was initially excluded from the YAP droplets, it later became localized to the rim of the droplets, as did newly transcribed RNA. This coincided in time with the expression of YAP downstream target genes in these cells. Therefore, YAP nuclear condensates dynamically restructure the overall genomic environment over time (Fig. 8g).

YAP constructs incapable of condensate formation (i.e., YAPΓTAD) could prevent both TEAD1 localization to clusters and YAP target gene expression during hyperosmotic stress. This suggested that the presence of YAP condensates during hyperosmotic stress is functionally relevant for YAP signaling. YAP’s TAD domain is rich in LCRs. Many proteins with LCRs undergo phase separation in response to stressors like arsenite^45,46^ or changes in salt concentration^33,47,48^. However, YAP formed condensates only in response to macromolecular crowders, such as sorbitol and PEG. This suggested that the specific structural features of YAP involved in its phase separation are poised for sensing macromolecular crowding. YAP’s paralog TAZ increased its tendency to form droplets under hyperosmotic stress (co-localizing with YAP in droplets). However, TAZ droplets were sometimes observed in the absence of hyperosmotic stress, as also reported in Wu et al.^33^. This suggested that differences in the properties of condensate formation by YAP and TAZ could underlie their differences in activation of downstream gene targets^49,50^. In fact, we found that TEAD1 clustering into condensates after hyperosmotic stress was significantly reduced in YAPΓTAD expressing cells, suggesting YAP acts upstream of TAZ in triggering the reorganization of TEAD1-based gene targets into clusters under these conditions. That said, further work will be needed to understand the relative roles of YAP and TAZ condensates in genome reorganization under different conditions^33^.

In summary, we’ve shown that YAP’s distinct localizations/functions in cytoplasm and nucleus can be controlled by its ability to undergo phase condensation in response to hyperosmotic stress. This allows YAP to be a sensor of mechanical forces in tissues that cause changes in macromolecular crowding. New strategies for modifying YAP target gene expression may thus become possible for altering the course of cancer and other diseases involving YAP signaling, for example, through droplet disrupters or reagents that change macromolecular crowding.

**Supplementary Fig. 1.**
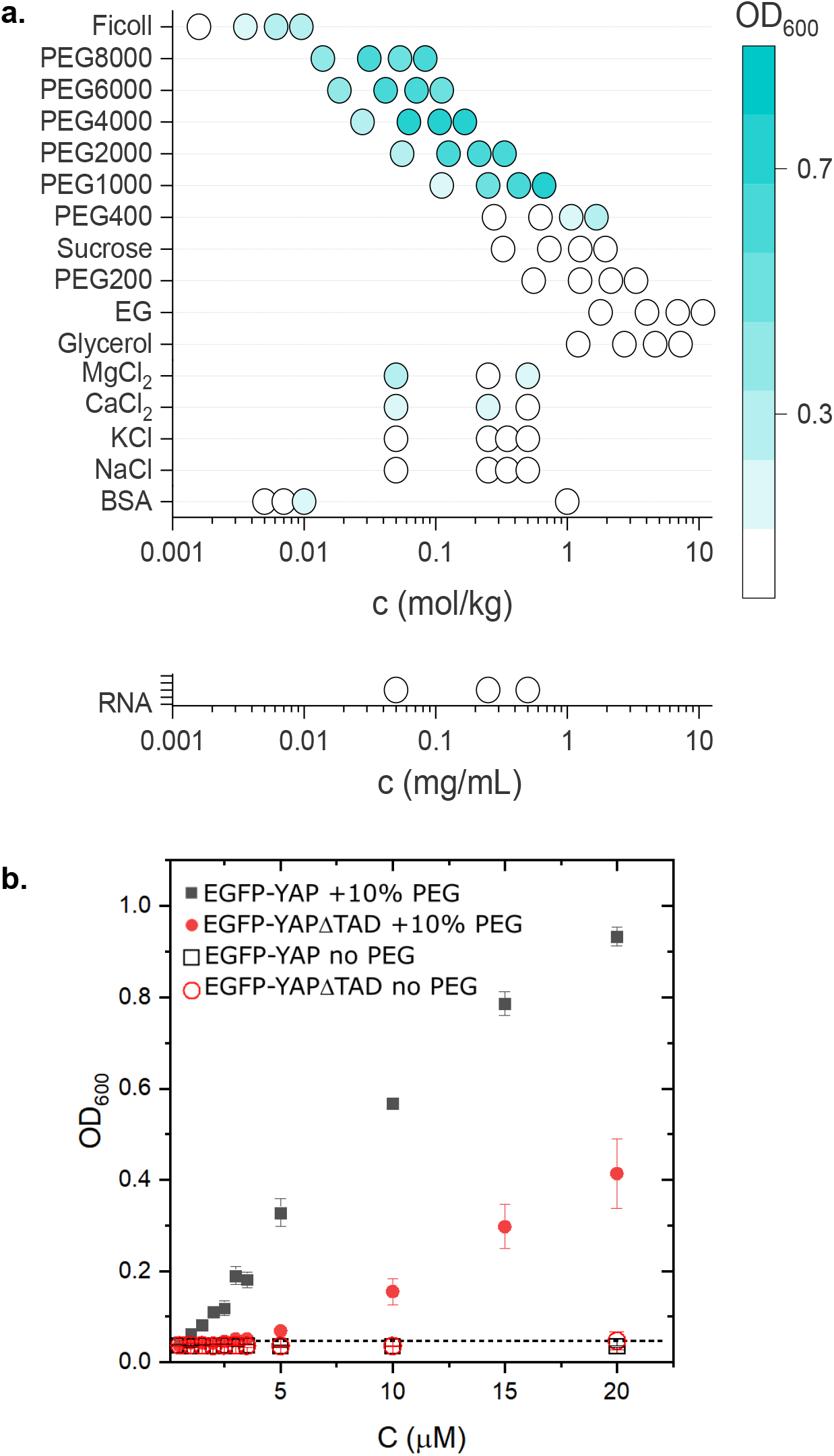
More characterizations of recombinant YAP protein in vitro. (a) Turbidity measurements of purified EGFP-YAP at different salt, BSA and RNA concentrations. (b) In vitro turbidity assay showing purified EGFP-YAP phase separates at much lower concentrations than EGFP-YAPΔTAD in the presence of 10% PEG2000. Error bars are standard deviations from three repeats.

**Supplemental Fig. 2.**
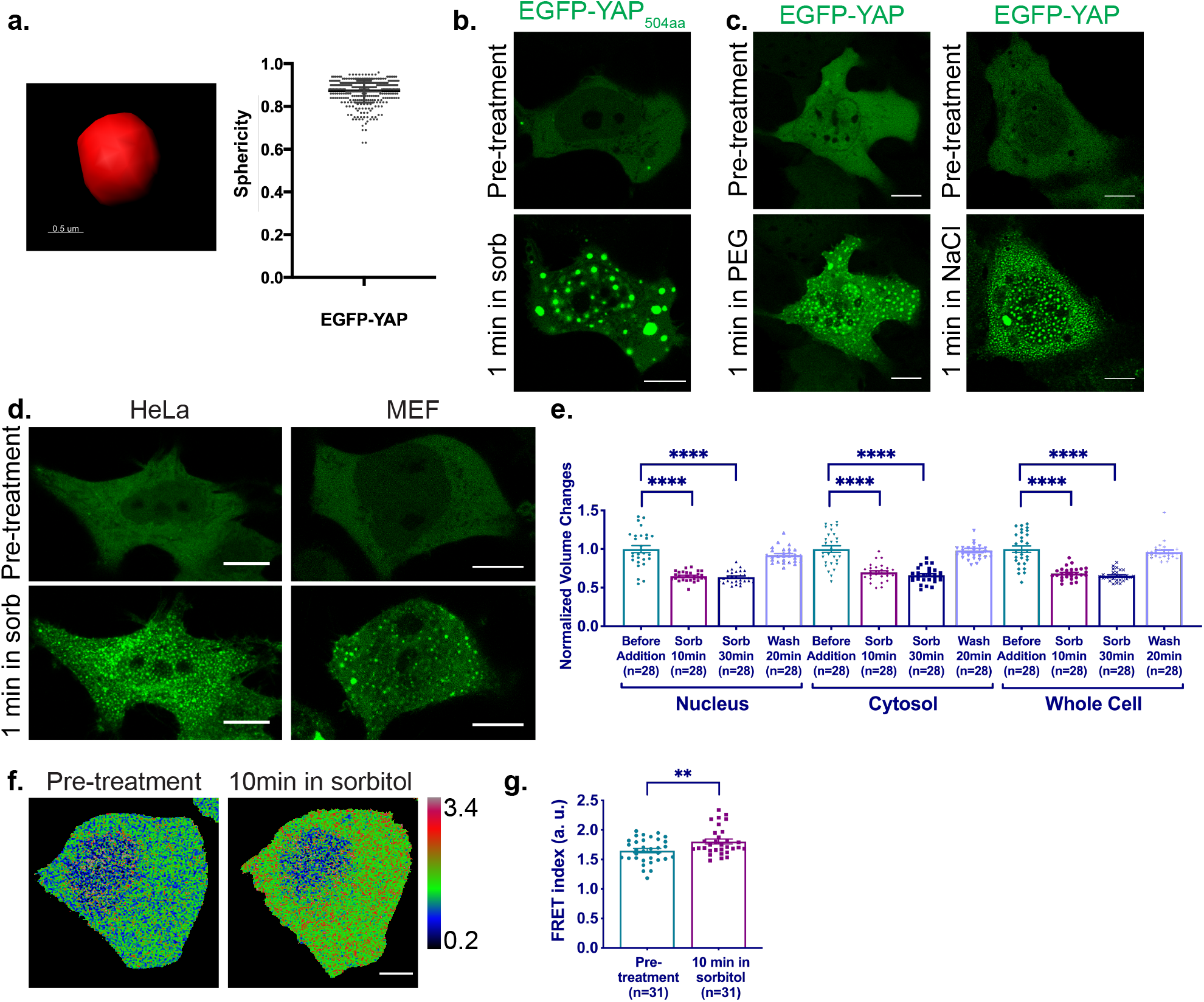
EGFP-YAP condensates form in hyperosmotic stress. Imaris 3-D rendering of an EGFP-YAP condensate, and quantification of sphericity of those condensates using Imaris. (b-d) Live-cell imaging of EGFP-YAP in HEK293T cells showing nuclear and cytoplasmic condensates are able to form with different isoforms of YAP (b) and in different hyperosmotic agents (c). (d) Live-cell imaging showing different cell types are able to form EGFP-YAP condensates under hyperosmotic stress. Scale bars are 10μm. (e) Quantification of normalized HEK293T nuclear, cytosolic and total volume before and after sorbitol treatment, and after wash. Unpaired t test. Comparing to volume before treatment. ****p<0.0001. Error bars show SEM. (f-g) Representative ratiometric images (f) and quantification (g) of crowding sensor FRET expressed in the same HEK293T cell before and after 0.2M sorbitol treatment. Rainbow RGB look-up table showing changes in FRET indices. Color bar: FRET index (a.u.). Paired t-test analysis. **p<0.01. Error bars show SEM.

**Supplemental Fig. 3.**
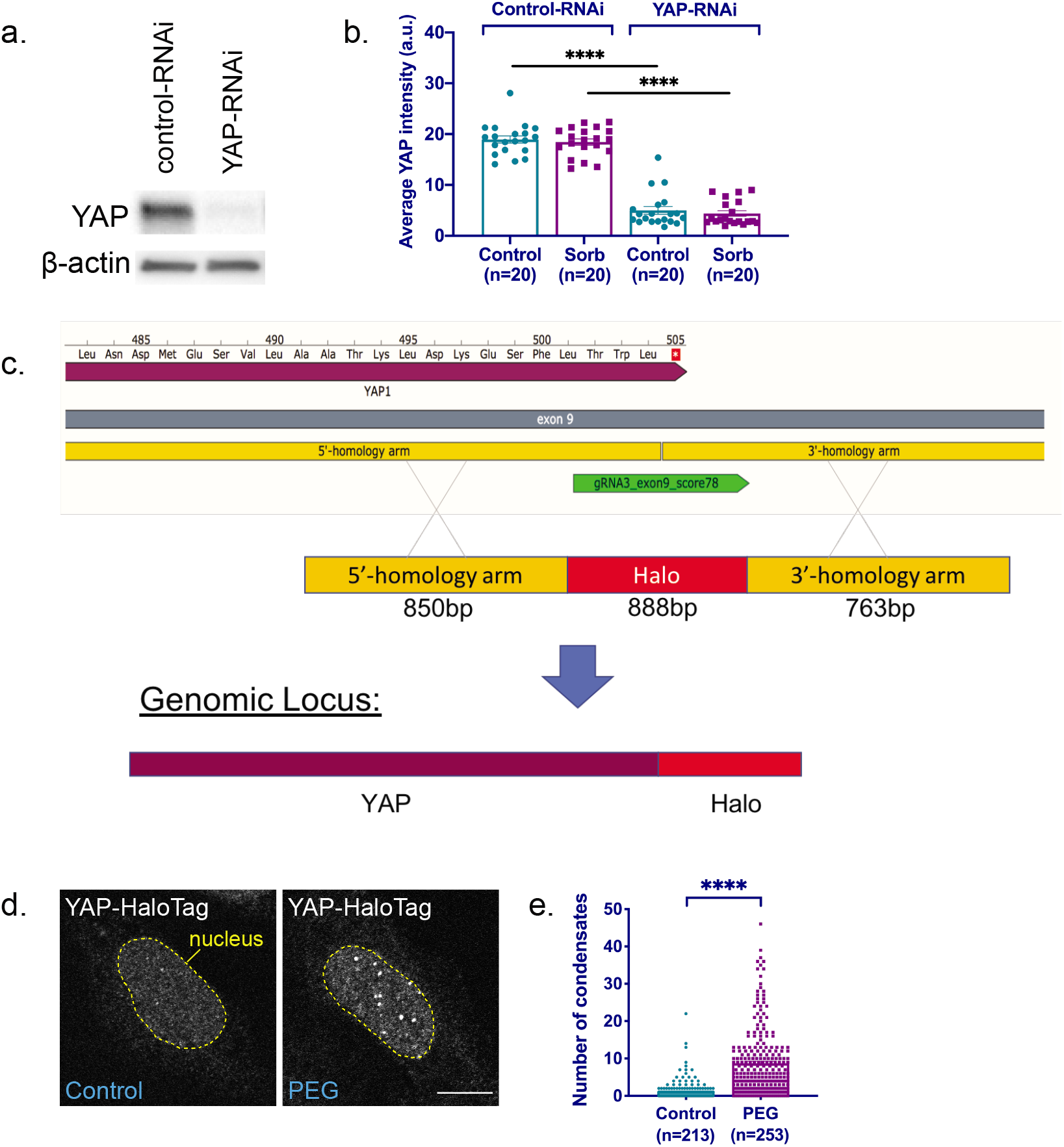
Endogenous YAP forms condensates. (a-b) Immunoblotting experiments (a) and quantifications of immunofluorescence YAP signal (b) indicate that YAP signal is effectively knocked down by YAP siRNA. Unpaired t-test is used in b. Error bars show SEM. ****p<0.0001. (c) Schematics of construction of CRISPR knock-in YAP-HaloTag U-2 OS cell line. Live-cell imaging (d) and quantification (e) showing nuclear YAP-HaloTag condensate labelled by JF549 Halo dye increases in number after PEG 300 treatment. Unpaired t-test analysis. Error bars show SEM. ****p<0.0001.

**Supplemental Fig. 4.**
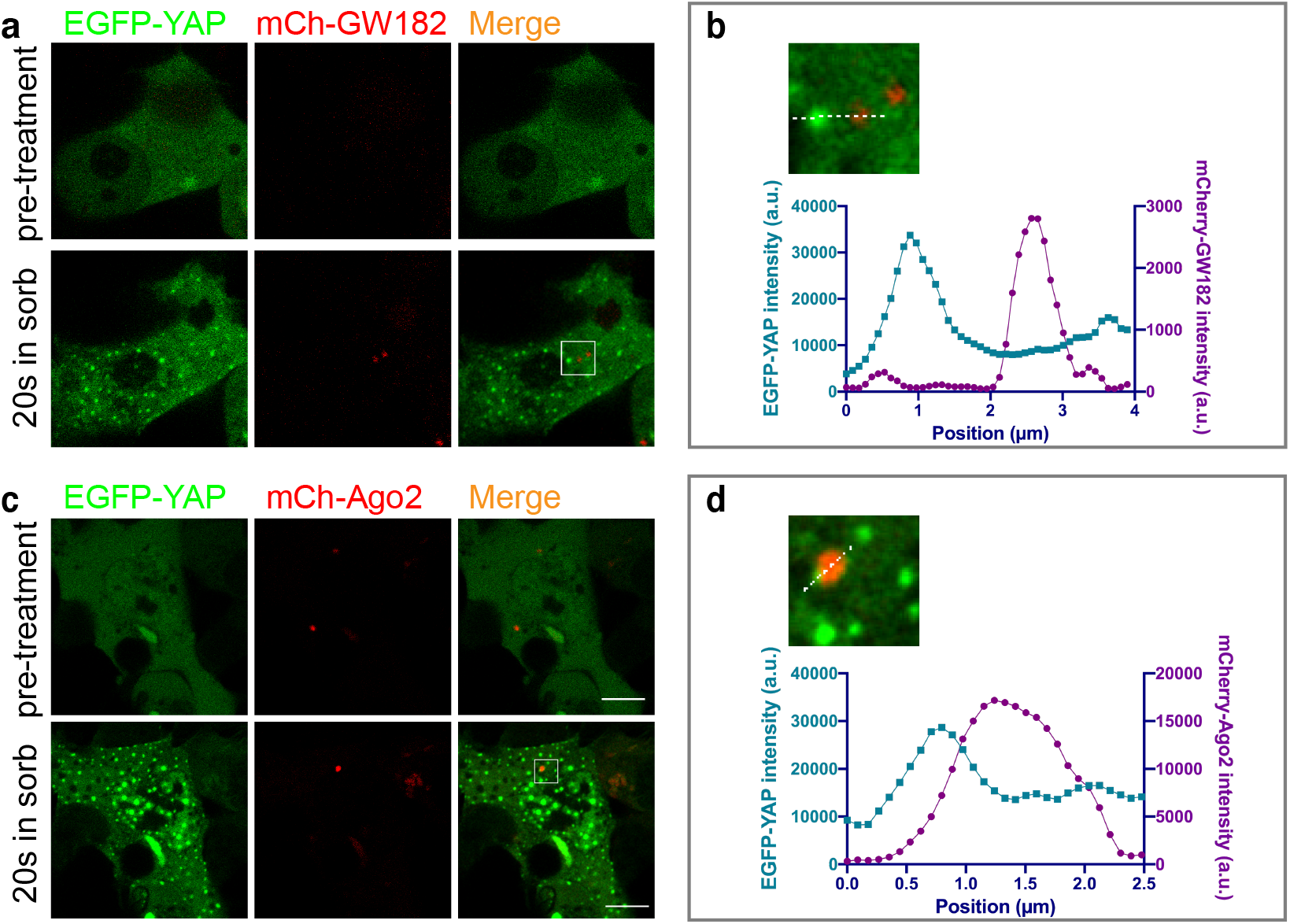
Characterization of cytoplasmic YAP condensates. (a) Livecell imaging showing no colocalization of cytoplasmic EGFP-YAP condensates with P-body component mCherry-GW182 after 0.2M sorbitol treatment for 20 sec in HEK293T cells. (b) Magnification of boxed region in (a) and line-scan. (c-d) Similar to (a-b), showing no colocalization of EGFP-YAP cytoplasmic condensates with mCherry-Ago2 after 0.2M sorbitol treatment for 20 sec. Scale bars are 10μm.

**Supplemental Fig. 5.**
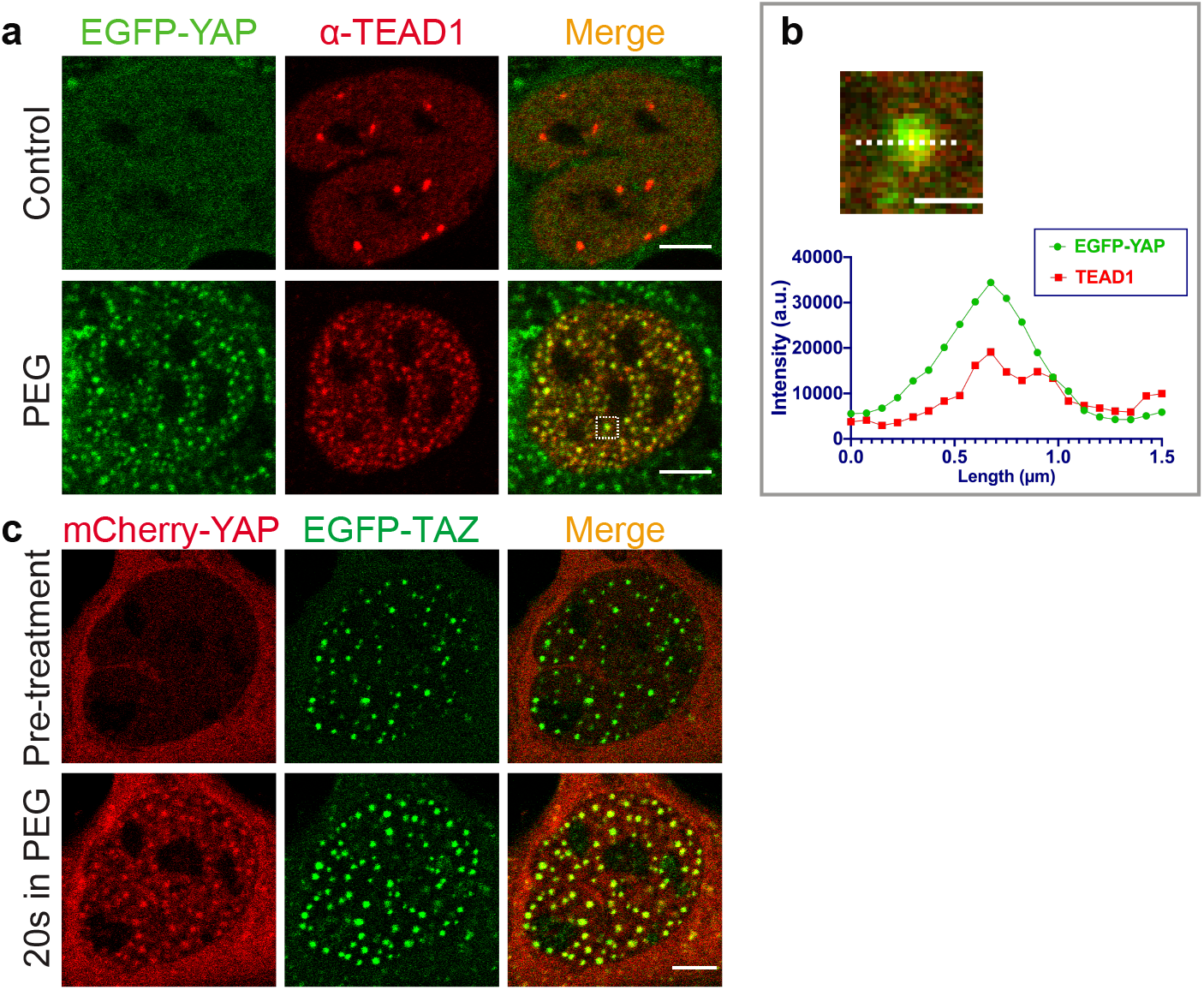
Characterization of nuclear YAP condensates. (a) Representative immunofluorescence images showing EGFP-YAP nuclear condensates colocalize with endogenous TEAD1 under hyperosmotic stress in U-2 OS cells. (b) Magnification of boxed region in a, and line scan. Scale bar: 1μm. (c) Live-cell imaging showing mCherry-YAP localizes to EGFP-TAZ condensates and new condensates after hyperosmotic stress in U-2 OS cells.

**Supplemental Fig. 7.**
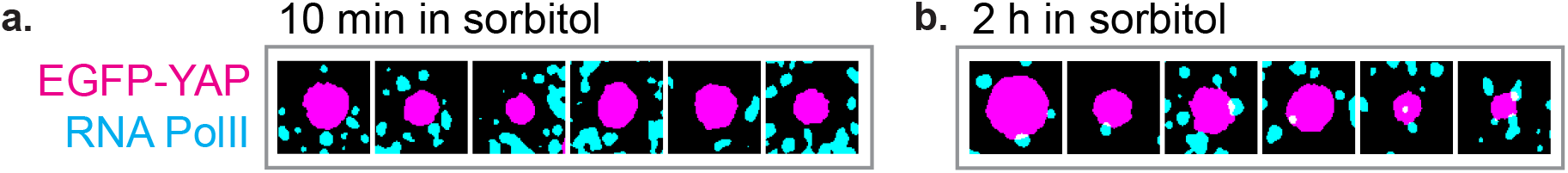
Dynamic localization of RNA Pol II relative to EGFP-YAP nuclear condensates after hyperosmotic shock. (a) More examples showing no colocalization of RNA Pol II (pSer2) with EGFP-YAP condensates at 10min in 0.2M sorbitol. Experiments are same as those in Fig. 7b, EGFP-YAP condensates are auto-thresholded and turned into mask using ImageJ, and pseudo-colored magenta. RNA Pol II immunofluorescence is autothresholded and turned into mask using ImageJ, and pseudo-colored cyan. (b) Similar to a, but showing more examples of enhanced localization of RNA Pol ll (pSer2) to the periphery of EGFP-YAP condensates at 2 h in sorbitol.

**Supplemental Fig. 8.**
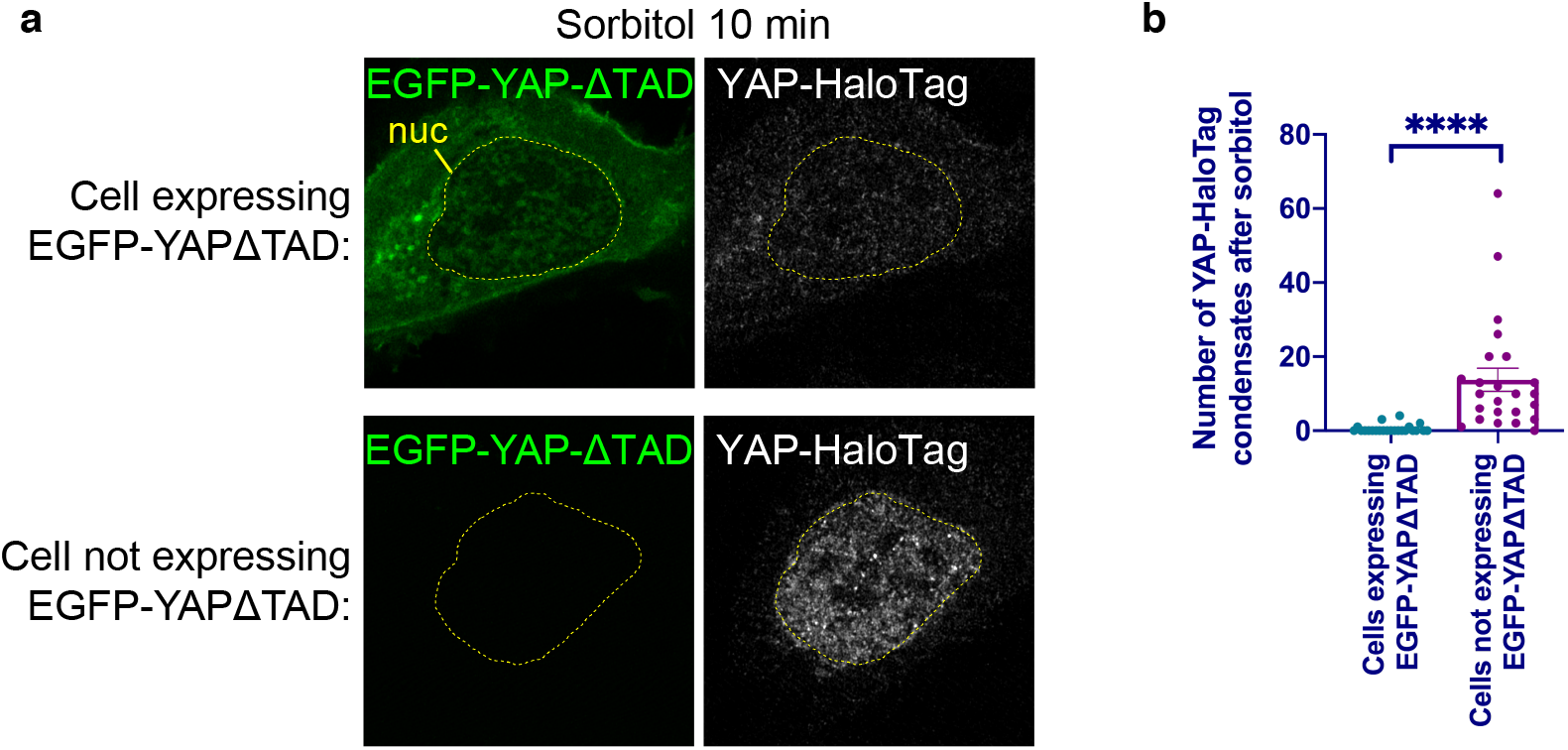
EGFP-YAPΓTAD mutant serves as a dominant negative protein that decreases endogenous YAP foci. (a) Immunofluorescence images of YAP-HaloTag U-2 OS cells 10 min in sorbitol treatment, with EGFP-YAPΔTAD overexpression (top row) or without (bottom row). (b) Quantification showing YAP-HaloTag U-2 OS cells have lower number of YAP-HaloTag endogenous YAP condensates after sorbitol treatment for 10 min, if they overexpress EGFP-YAPΓTAD construct. Unpaired t-test is conducted. ****p<0.0001. Error bars show SEM.

