## Supplementary files 1-3 for "Phase separation of YAP reorganizes genome topology for long-term, YAP target gene expression"

**Supplementary Material #1: Material and Methods**

Constructs

pEGFP-C3-hYAP1 (Addgene #17843) and pEGFP C3-Mst2 (Addgene #19056) were gifts from Marius Sudol. mCherry-Dcp1a was a gift from Pick-Wei Lau. mCherry-NLK construction: PCR-amplify NLK from pDONR223-NLK (gift from William Hahn & David Root, Addgene #23642) with 5'- ctcaagcttcgaattctgcaCTTCCACACCTCCCTCCTC-3' and 5'- gatccggtggatcccgggccCTCCCACACCAGAGGAGATG-3'; PCR-amplify pmCherry-C1 (Clonetech) with 5'- GGCCCGGGATCCACCGGA-3' and 5'-TGCAGAATTCGAAGCTTGAGCTC-3'. Assemble amplified NLK and mCherry-C1 with NEBuilder HiFi DNA Assembly Master Mix. pET28b-EGFP-YAP construction: pET28b (+) (EMDMillipore) and pEGFP-C3-hYAP1 are each double-digested with NheI and EcoRI (NEB) and ligated together by T4 DNA Ligase. Subsequently, EGFP-YAP is brought in-frame with 6xHis-thrombin by QuikChange Lightning Site-Directed Mutagenesis Kit (Agilent 210519) by primers: 5'- gctaccggtcgccacatggtgagcaagg-3' and 5'-ccttgctcaccatgtggcgaccggtagc-3'. For generation of YAP truncations, pEGFP-C3-hYAP1 (addgene # 17843) was used as template to amplify YAP-ΔP (FWR 5'-GGAAGATCTTCC CAT CAG ATC GTG CAC GTC CG-3' and REV 5'-CCGGAATTCCGG CTA TAA CCA TGT AAG AAA GCT TTC TTT ATC -3') and YAP-ΔTAD (FWR 5'- GGAAGATCTTCC ATG GAT CCC GGG CAG CAG -3' and REV 5'-CCGGAATTCCGG CTA ACC CAT GAC GCC TCC CTG -3') by PCR and then these were cloned into Bgl II/EcoRI sites of pEGFP-C3. To generate pEGFP-YAP-S127A, pEGFP-YAP-S128A constructs, pEGFP-C3-hYAP1 construct is used as template, and QuikChange Lightning Site-Directed Mutagenesis Kit is used to make point mutations using primers: 5'-gagaagctggagaggcatgagctcgaacatgct-3' (forward) and 5'-agcatgttcgagctcatgcctctccagcttctc-3' (reverse) for S127A, and 5'-gttcgagctcattccgctccagcttctctgcAGT-3' (forward), 5'-gcagagaagctggagcggaatgagctcgaacATGCT-3' (reverse) for S128A.

In vitro EGFP-YAP expression, protein purification, and phase separation experiments

pET28b-EGFP-YAP was used to transform E.Coli BL21 DE3 RIPL cells (Invitrogen) using standard supplier protocol. Cells were grown to OD 0.6, lyzed using a tip sonicator, and purified using His-tag affinity chromatography on a GE-Healthcare AKTA system. The purified protein was concentrated to ~ 1 mg/mL and run on a Sephadex 200 size exclusion column in 20 mM Tris pH 8.0. The collected fractions contained the clean protein, as verified with SDS PAGE and MALDI-TOF mass spectrometry. Phase separation conditions were assessed by sample turbidity, measured through the OD at 600 nm of EGFP-YAP solutions in 96-well clear-plastic plates using a Molecular Designs SpectraMax plate reader. To determine the partition coefficient, $K_P$ of YAP (Fig. 3I) between solution and protein-rich droplets, phase separation reactions were prepared in 1.5 mL tubes with 40 μM EGFP-YAP, and different PEG sizes in concentrations ranging from 0 to 35 wt% (Sigma), and 20 mM Tris buffer pH 8.0. At given times, the reaction solution was gently stirred, and a 50 μL aliquot was removed. The aliquot was spun down at max speed for 15 minutes. A bright green pellet was observed on the bottom of the tubes, and the supernatant was carefully removed, diluted 5x in tris buffer, and placed in a 96-well plate for absorbance measurements at the maximal absorbance wavelength of EGFP, 488 nm. The concentration of EGFP-YAP in the soluble phase was estimated based on EGFP absorbance.

Confocal imaging of purified EGFP-YAP was done on an LSM-880 equipped with an Ar laser line. Imaging chambers were prepared on clean glass slides using a 120 µm double-sided sticker (Grace Biolabs), and sealed with a 1.5 coverslip. Imaging was done on a single slice, with the pinhole opened to 2 Airy units.

Cell Culture, Transfection, siRNA and Live-cell imaging
U2-OS and HEK293T cells are cultured in complete medium: Dulbecco's modified Eagle's medium (DMEM) supplemented with 10% fetal bovine serum (FBS) (Gibco), 100 units/ml penicillin/ streptomycin (Corning), and 2mM L-glutamine (Corning) at 37°C and 5% CO2. For confocal imaging, cells are cultured on 8-well LabTek chambered coverglass dishes (Thermo Scientific), and transfected with Lipofectamine 3000 Reagent (Thermo Fisher). Cells are imaged 18hrs after transfection on a Zeiss LSM780 confocal microscope at room temperature with Plan-Apochromat 63x/1.40 Oil objective. For siRNA experiments, YAP siRNA (Thermo Fisher Silencer Select s20367) or negative control (Thermo Fisher AM4611) is transfected into cells (pre-seeded on 6-well plate) using Lipofectamine™ RNAiMAX Transfection Reagent (Thermo Fisher 13778075). 48 hr after transfection, cells are replated onto 6-well plates for confirming knock down by immunoblotting, or to 8-well LabTek chambered coverglass dishes for imaging on the following day. Images are taken every 30 sec, in a z-stack of 3 slices with an interval of 0.4µm. For volume measurements, cells pre-transfected with EGFP-YAP are labelled with Hoechst 33342 (Thermo Fisher 62249) for 30 min prior to confocal imaging. Dual color imaging and z-sectioning is done across the whole cell and nuclear volume, with an interval of 0.5µm. Cell and nuclear volume measurements are done with Imaris software (Bitplane) using surface reconstruction tool, and cytoplasmic volume is calculated by substracting cell volume (EGFP channel) with nuclear volume (Hoechst channel). To visualize fusion of YAP condensates, images are taken every 4.5 sec with Definite Focus. Zoomed-up images are smoothened once using ImageJ for display.

Fluorescence Recovery after Photobleaching (FRAP)
HEK293T cells expressing EGFP-YAP is treated with 0.2M sorbitol, and FRAP experiments are done on cytoplasmic or nuclear YAP condensates formed immediately after treatment using Bleaching mode in Zen. Region of interest is selected on either the entire condensate or part of it using a rectangular box of approximately 1µm x 1µm in size. 20 iterations of bleaching are done with 100% 488nm Argon laser. 5 rounds of imaging are done prior to, and 300 rounds of imaging are done after bleaching until fluorescence signal plateau, with an interval of 450 msec. $t_{1/2}$ is calculated with FRAP module in Zen software, using fit formula for 1 diffusion component.

RT-PCR
HEK293T cells are serum-starved for 1hr, and subject to control serum-free medium or 0.2M sorbitol serum-free medium, according to methods described in Hong et. al[1]. Total RNA is isolated from HEK293T cells using Qiagen RNeasy mini kit (Qiagen 74104) and

converted to cDNA with Thermo Fisher High-Capacity cDNA Reverse Transcription Kit (Thermo Fisher 4368814). RT-PCR is carried on an Agilent AriaMx 96 Real-Time PCR System, with Platinum SYBR Green qPCR Supermix-UDG (Thermo Fisher 11733038). The following primers are used: Gapdh: 5'-CTCCTGCACCACCAACTGCT-3' (forward), 5'-GGGCCATCCACAGTCTTCTG-3' (reverse). Ctgf: 5'-AGGAGTGGGTGTGTGACGA-3' (forward), 5'-CCAGGCAGTTGGCTCTAATC-3' (reverse). Cyr61: 5'-CCTCGGCTGGTCAAAGTTAC-3' (forward), 5'-TTTCTCGTCAACTCCACCTC-3' (reverse). mRNA levels are normalized to that of Gapdh.

Generation of YAP-HaloTag CRISPR knock-in U-2 OS cell line

Singe-guide RNA (sgRNA) targeting +/- 100bps around the stop codon of Yap gene was designed using the web-based CRISPR design tool (http://crispr.mit.edu). One guide is chosen due to its spanning the stop codon and high score: tacatggttatagagccctc. DNA oligonucleotides with BbsI restriction sites are ordered from Integrated DNA Technologies (IDT). pSpCas9(BB)-2A-GFP (PX458) vector (Addgene 48138, a gift from Feng Zhang)[2] was digested using BbsI and ligated with annealed gRNA fragment to form Yap gRNA-Cas9 plasmid.

The homology repair fragment spanning Yap stop codon (~800bp on each side) and containing Halo protein-coding gene is synthesized with gBlock IDT (see supplementary material 2 for sequence). pUC57-mini vector (GenScript) is linearized with EcoRV restriction enzyme, and ligated with Halo homology repair fragment using NEBuilder® HiFi DNA Assembly Master Mix (NEB E2621). All plasmids are sequence-verified by Eurofins Genomics.

Yap gRNA-Cas9 and Halo homology repair constructs are co-transfected into U-2 OS cells with Lipofectamine 3000 Transfection Reagent (Thermo Fisher L3000015). 48 h after transfection, GFP-positive cells are selected using fluorescence-activated cell sorting (FACS, National Eye Institute Flow Cytometry Core). Cells are grown for an additional week, and stained by 100nM JF549 Halo dye (gift of Luke Lavis, HHMI Janelia Research Campus) for 30 min, washed 3 times with 1xPBS, and FACS-sorted for a second time for Halo dye-positive and GFP-negative cells. Individual cells are grown in single wells of a 96-well dish supplemented with 20% FBS DMEM medium, and waited for 1 week for positive clones to form. Cell clones are seen in 40% of the wells. Correct genome insertion is verified by genome DNA extraction (QuickExtract™ DNA Extraction Solution, Lucigen QE09050), and PCR using primers spanning homology arms and Halo tag: 5'-attcctgggacaaatgtggaccttg-3' (forward), 5'-tcaggtctggagcaatgcagcgatg-3' (reverse); and 5'-tgaatcctgttgaccgcgagccactg-3' (forward), 5'-tagaattcagtctgcctgagggctc-3' (reverse). Increased protein size of YAP marked by Halo is further confirmed by immunoblotting with anti-YAP antibody (Cell Signaling 14074S).

Live-cell imaging of U-2 OS YAP-HaloTag cell line

YAP-HaloTag U-2 OS cells are pre-plated on 8-well LabTek chambered coverglass dishes for at least 16 hrs in complete medium. Prior to imaging, incubate cells with complete medium containing 100 nM JF549 Halo dye for 30 min, and then wash 3 times with 1x PBS medium before changing back to complete medium. Complete medium and 1x PBS need to be isotonic (no old medium with water evaporated) to prevent early formation of YAP-HaloTag condensates. Imaging is done on a Zeiss LSM880 scope with Airyscan (Superresolution mode), using a Plan-Apochromat 63x/1.4 oil DIC M27 objective. RFP 561nm laser is used to excite the Halo dye, and is kept under 0.7% power to minimize photobleaching. Pinhole is kept at 132 μm, and pixel sizes are 35 nm x 35 nm.

Kidney dissection and Immunofluorescence

Kidney tissue from 2 WT mice were immersion-fixed with 4% PFA in phosphate buffered saline (PBS) for 48 hr at 4 °C. For immunofluorescence microscopy (IF) stainings, fixed kidney tissue was mounted in 5 % low melting agarose in PBS and sectioned into 75 μm thick slices using a vibratome (Leica). Floating sections were permeabilized in 0.5 % Triton-X100 in PBS for 1 hr, and blocked by incubation with blocking buffer (20% FBS + 0.2 % Triton-X100 in PBS) for 2 hr. Sections were incubated sequentially with a primary antibody against YAP (1:50) (Cell Signaling) in blocking buffer for 72hrs, washed 6 times for 10 min with 0.2 % Triton-X100 in PBS, incubated with secondary antibody labelled with Alexa fluorophore 488 (1:500), Hoechst (1:5000) and Alexa fluorophore 568-conjugated Phalloidin (1:150) in blocking buffer overnight, and washed again with 0.2% Triton-X100 in PBS 6 times for 10 min. Stained sections were incubated in 50% glycerol overnight, and then mounted in slides in 80% glycerol. Fixed kidney samples were imaged with a Leica SP8 laser scanning confocal microscope (DMI8-CS) using a 63x 1.4 oil objective, 405, 488 and 552 laser lines and HyD detectors. Images of the kidney cortex and medulla were acquired.

Immunofluorescence and Nascent RNA Pulse Labeling

HEK293T cells are plated on coverslips precoated with fibronectin (7.5μg/ml, Millipore, FC010) for 16 h, before being fixed with 4% paraformaldehyde (PFA, EMS), permeabilized with 0.5% TritonX-100, and blocked with 3% BSA in 1xPBS. Incubate with primary antibodies in 1% BSA overnight at 4°C, and then incubate with Alexa Fluor-conjugated secondary antibodies. The following primary antibodies are used: anti-YAP (Cell Signaling 14074S); anti-TEAD1 (BD Biosciences 610922); anti-RNA polymerase II CTD repeat YSPTSPS (phospho S2) (Abcam ab193468); anti-G3BP1 (proteintech 13057-2-AP); anti-PML (Abcam ab96051); and anti-Coilin (Abcam ab87913). Nascent RNA is labelled using Click-iT™ RNA Alexa Fluor™ 594 Imaging Kit (Thermo Fisher C10330). All samples are transiently labelled with 2.5mM 5-ethynyl uridine (EU) for 5 min prior to fixation with 4% PFA. Specifically, sample treated with 5 min sorbitol is supplemented with 0.2M sorbitol with 2.5mM EU for 5 min. Sample treated with 2 h sorbitol is supplemented with 0.2M sorbitol for 1 h 55 min before changing to 0.2M sorbitol supplemented with 2.5mM EU and incubated for 5 min. Each sample is incubated with GFP-Booster_Atto488 (ChromoTek gba488-100) after fixation and permeabilization, before performing Click chemistry to visualize EU following the manufacturer instructions. For imaging and quantification, at least 15 fields of view per coverslip are randomly chosen by Hoechst nuclear staining and imaged by Zeiss LSM780, LSM880 confocal microscope or Airysan. At least 3 different coverslips are quantified per treatment type.

Colocalization

Colocalization of two channels are done with ImageJ Coloc 2 plugin. ROI is chosen on the either the cytoplasm or the nucleus. Pearson's R value is used for measuring colocalization of two channels.

**3D ATAC-PALM imaging:**

Sample preparation for 3D ATAC-PALM imaging

We prepared samples for 3D ATAC-PALM experiments as reported previously[3,4]. HEK293 cells were plated onto 5mm coverslips (Warner Instruments, cat#64-0700) at around 80% confluency with proper coating one day before experiment. Control cells or cells under 5 min sorbitol treatment were fixed with 4% paraformaldehyde (Electron Microscopy Sciences, Cat# 15710) for 10 min at room temperature. After fixation, cells were washed three times in 1×PBS for 5 minutes and then permeabilized in ATAC lysis buffer (10 mM Tris–Cl, pH 7.4, 10 mM NaCl, 3 mM MgCl2, 0.1% Igepal CA-630) for 10 min at room temperature. After permeabilization, the coverslips were washed twice in 1×PBS. The transposase mixture solution was prepared according to previous research [3] and was added to the cells. The sample was incubated in a humidity chamber box for 30 min at 37 °C. After incubation, the coverslips were washed three times with 1×PBS containing 0.01% SDS and 50 mM EDTA for 15 min at 55 °C before mounted onto the Lattice light-sheet microscope (LLSM) slot for 3D ATAC-PALM imaging.

3D ATAC-PALM image acquisition and processing

The 3D ATAC-PALM imaging was performed following published procedures[3] by using the lattice light-sheet microscopy[5]. The light sheet was generated from the interference of highly parallel beams in a square lattice and dithered to create a uniform excitation sheet. The inner and outer numerical apertures of the excitation sheet were set to be 0.44 and 0.55 during experiments, respectively. In order to maintain stable imaging conditions in particular constant salt concentration, a Variable-Flow Peristaltic Pump (Thermo Fisher Scientific) was used to connect a 2L reservoir with the imaging chamber with 1×PBS circulating through at a constant flow rate. Labelled cells were placed into the imaging chamber and each image volume includes 100~200 image frames. Initially, fluorescent PA-$JF_{549}$ dye were pushed into the dark state through repeated photo-bleaching by maximal laser power (2W) (MPB Communications Inc., Canada). The samples were then imaged by iteratively photo-activating each focal plane with very weak intensity 405 nm light (<0.05 mW power at the rear aperture of the excitation objective and 6W/$cm^2$ power at the sample) for 8 ms, followed by exciting each plane with a 2W 560 nm laser at its full power (26 mW power at the rear aperture of the excitation objective and 3466 W/$cm^2$ power at the sample) for 20 ms exposure time. The specimen was illuminated when laser light went through a custom 0.65 NA excitation objective (Special Optics, Wharton, NJ) and the fluorescence generated within the specimen was collected by a detection objective (CFI Apo LWD 25×W, 1.1 NA, Nikon), filtered through a 440/521/607/700 nm BrightLine quad-band bandpass filter (Semrock) and N-BK7 Mounted Plano-Convex

Round cylindrical lens (f = 1000 mm, Ø 1", Thorlabs), and eventually recorded by an ORCA-Flash 4.0 sCMOS camera (Hamamatsu).

We embedded nano-gold fiducials within the coverslips for drift correction during imaging process as previously described[6]. ATAC-PALM Images were taken to construct a 3D volume when the sample was moving along the sample axis. Individual volumes per acquisition were automatically stored as Tiff stacks, which were then analyzed by in-house scripts written in Matlab. The cylindrical lens introduced astigmatism in the detection path and recorded each isolated single molecule with its ellipticity, thereby encoding the 3D position of each molecule relative to the microscope focal plane. The localization precision was estimated to be 26±3 nm and 53±5 nm for xy and z respectively by calculating the standard deviation of all the localizations coordinates (x, y and z) after the nano-gold fiducial correction.

Raw 3D ATAC-PALM images were processed following previous published procedures Specifically, a background image with gray values taken without illumination was subtracted from each frame of the current raw image. Photo-activated images were then filtered by subtracting Gaussian filtered images at two different standards:

$$\mathrm{I}_{filtered} = \mathrm{I}_{raw} \otimes K_{low} - \mathrm{I}_{raw} \otimes K_{high} \qquad \textbf{(Equation S1)}$$

Where $\otimes$ is the convolution operator, and $K_{low}$ and $K_{high}$ represent 5×5 pixel two-dimensional Gaussian function with standard deviation $\sigma$=2 (low) and 1 (high):

$$K = 2\pi^{-3/2}\sigma^{-2} \cdot e^{-\frac{(x-3)^2}{2\sigma^2}-\frac{(y-3)^2}{2\sigma^2}} \qquad \textbf{(Equation S2)}$$

The local maxima were then determined on these filtered images and the coordinates were estimated and used as the true position of the isolated single molecule.

In order to precisely determine the z position of each isolated localization, cylindrical lens were used and the corresponding astigmatic PSF function could be formulated as a Gaussian function with separate x and y deviations. Importantly, these deviations encoded the relative z-offset of the signal with respect to the focal plane.

$$PSF(x,y,z) = \frac{1}{2\pi\sigma_x\sigma_y} \cdot e^{-\left(\frac{x^2}{2\sigma_x{}^2}+\frac{y^2}{2\sigma_y{}^2}\right)} \qquad \textbf{(Equation S3)}$$

Where

$$\sigma_x = c_1 + c_2 z + c_3 z^2$$

$$\sigma_y = c_4 + c_5 z + c_6 z^2$$

Here $\sigma_x$ and $\sigma_y$ were polynomial function of z. The constants $c_1$~$c_6$ can be fitted by scanning bright and regular beads through the focal plane using piezoelectric stage.

To derive the ATAC-PALM intensity map in order to compare with GFP-YAP signal, ATAC-PALM localizations were binned within a cubic of 100 nm with a 3D Gaussian filter and a convolution kernel of 3×3×3.

3D pair auto-correlation function

As described previously[7], the pair correlation function $g_0(r)$ or radial distribution function measures the probability $P$ of finding a localization of accessible chromatin in a volume element $dV$ at a separation $r$ from another accessible chromatin site:

$$P = \rho \cdot g_0(r) \cdot dV \quad \textbf{(Equation S4)}$$

Where ρ represents the mean density of accessible chromatin in the nucleus.
3D $g_0(r)$ was computed by

$$g_0(r) = \left[ \frac{V}{N-1} \cdot \frac{3}{4\pi(3r^2 \cdot \Delta r + 3r \cdot \Delta r^2 + \Delta r^3)} \right] \cdot \left[ \frac{1}{N} \sum_{i=1}^{N} \sum_{i \neq j} \delta(r - r_{ij}) \right] \quad \textbf{(Equation S5)}$$

N is the total number of localizations and (N-1)/V is the average localization density within the 3D nuclear volume *V*. $\Delta r = 50$ nm is the binning width used in the analysis. The item $4\pi(3r^2 \cdot \Delta r + 3r \cdot \Delta r^2 + \Delta r^3)/3$ represents the shell-shape volume between the search radius from *r* to $r + \Delta r$. The Dirac Delta function is defined by

$$\delta(r - r_{ij}) = \begin{cases} 1 & r - r_{ij} \leq \Delta r \\ 0 & r - r_{ij} > \Delta r \end{cases}$$

Where $r_{ij}$ represents the pair-wise Euclidean distance between localization point $i$ and $j$. To eliminate the boundary effect in calculation of $g_0(r)$ from a finite 3D volume, we generated $g_r(r)$ from uniform distributions with the same localization density in the same volume as real data using the Matlab convex hull function. Accordingly, the final normalized 3D pair auto-correlation function $G(r)$ was calculated by:

$$G(r) = \frac{g_0(r)}{g_r(r)} \quad \textbf{(Equation S6)}$$

Whereas $G(r)$ can be used as *bona fide* criteria for estimating the clustering effect of a group of spatial localizations, it has to be adjusted regarding the inevitable localization uncertainty of the PSF. If the distribution is purely random the revised pair auto-correlation $G_R(r)$ considering PSF uncertainty is given by

$$G_R(r) = G_{PSF}(r) + 1 \quad \textbf{(Equation S7)}$$

with

$$G_{PSF}(r) = \frac{1}{8\pi^{3/2}\sigma^3} e^{-r^2/4\sigma^2} \quad \textbf{(Equation S8)}$$

where *ρ* represents the average surface density of molecules, $G_{PSF}(r)$ denotes the correlation of uncertainty in the PSF and *σ* for standard deviation. The spatial autocorrelation of accessible chromatin can be approximated by an exponential function:

$$g(r) = A \cdot e^{-r/\varepsilon} + 1 \quad \textbf{(Equation S9)}$$

where *ε* and A denotes the correlation length and amplitude of the accessible chromatin cluster, respectively. The total correlation can be described by

$$G(r) = G_{PSF}(r) + \left( A \cdot e^{-r/\varepsilon} + 1 \right) \otimes G_{PSF}(r) \quad \textbf{(Equation S10)}$$

The raw data were fitted by using Equation S10 to derive $g(r)$ and parameters *A* and *ε*. The approximated function $g(r)$ was used throughout the paper for pair auto-correlation analysis. Curve fitting was performed using the trust-region method implemented in the Curve Fitting Matlab toolbox.

DBSCAN analysis

The DBSCAN algorithm (Density-Based Spatial Clustering of Applications with Noise) was adopted to map and visualize individual local ACDs (core DBSCAN Matlab code from http://yarpiz.com/255/ypml110-dbscan-clustering). The algorithm first finds neighboring data points within a sphere of radius *r*, and adds them into same group. In parallel, a predefined threshold minimal points (*minPts*) was used by the algorithm to justify whether any counted group is a cluster. If the number of points within a group is less than the threshold *minPts*, the data point is classified as noise. We implemented DBSCAN analysis on the localization data for both the control and 5min sorbitol-treated cells by using 150 nm as the searching radius (*r*) (peak radius from the Ripley's H function analysis) and empirically setting *minPts* as 10. To reconstruct the *iso*-surface for each identified accessible chromatin cluster, the convex hull of each accessible chromatin cluster was calculated and visualized by using Matlab. The volume of the convex hull was computed, and the normalized cluster radius (calculated from a sphere with equal volume) was calculated and shown in violin plot.

**Supplementary Material #2: gBlock sequence of Yap homology arms with HaloTag-coding sequence.**

ccacctgacgtcccaatgatgaagctctgaggagtgttaatgttgtcatatgcaaaaaggctctgtggccaagtaagt
ttgggaaacctagaagcaaggctaggaagatggatttattcaaagccctctgggaaggcttgttttgctgttaggaagg
tgcatctcctaagggtagaataaagtattcttgcagtctttctattttagtgtttcccaaagtatatttaaggccaaggtaggt
ctataccactcatgtttatgtttagtcttattatggtggtgcttatctggaagacagggtagtgatgctgtcgtgttgtgtacttg
ttctttctcccgctgctgcatataatttttcaagggctttttattttttcatgactcctttagtgctcagcacataatagcaatca
agctgttattggatatctgaacaatgttgaagcaaaagtaaacttggcattgttattttgttatctgtgagttgtttgacctgat
ataaattctaggtatcagtttaggtcagtcaggagcgcttttctttccctgtgcatttctctgtgctcatatgatacagccctg
atgttagcttttcaaaaaggactttgttatctctgtgtgtttccactaggtgatactatcaaccaaagcaccctgccctcaca
gcagaaccgtttcccagactaccttgaagccattcctgggacaaatgtggaccttggaacactggaaggagatgga
atgaacatagaaggagaggagctgatgccaagtctgcaggaagctttgagttctgacatccttaatgacatggagtct
gttttggctgccaccaagctagataaagaaagctttcttacatggttaGCAGAAATCGGTACTGGCTTTC
CATTCGACCCCCATTATGTGGAAGTCCTGGGCGAGCGCATGCACTACGTCGAT
GTTGGTCCGCGCGATGGCACCCCTGTGCTGTTCCTGCACGGTAACCCGACCTC
CTCCTACGTGTGGCGCAACATCATCCCGCATGTTGCACCGACCCATCGCTGCA
TTGCTCCAGACCTGATCGGTATGGGCAAATCCGACAAACCAGACCTGGGTTAT
TTCTTCGACGACCACGTCCGCTTCATGGATGCCTTCATCGAAGCCCTGGGTCT
GGAAGAGGTCGTCCTGGTCATTCACGACTGGGGCTCCGCTCTGGGTTTCCACT
GGGCCAAGCGCAATCCAGAGCGCGTCAAAGGTATTGCATTTATGGAGTTCATC
CGCCCTATCCCGACCTGGGACGAATGGCCAGAATTTGCCCGCGAGACCTTCCA
GGCCTTCCGCACCACCGACGTCGGCCGCAAGCTGATCATCGATCAGAACGTTT
TTATCGAGGGTACGCTGCCGATGGGTGTCGTCCGCCCGCTGACTGAAGTCGA
GATGGACCATTACCGCGAGCCGTTCCTGAATCCTGTTGACCGCGAGCCACTGT
GGCGCTTCCCAAACGAGCTGCCAATCGCCGGTGAGCCAGCGAACATCGTCGC
GCTGGTCGAAGAATACATGGACTGGCTGCACCAGTCCCCTGTCCCGAAGCTGC
TGTTCTGGGGCACCCCAGGCGTTCTGATCCCACCGGCCGAAGCCGCTCGCCT
GGCCAAAAGCCTGCCTAACTGCAAGGCTGTGGACATCGGCCCGGGTCTGAAT
CTGCTGCAAGAAGACAACCCGGACCTGATCGGCAGCGAGATCGCGCGCTGGC
TGTCGACGCTCGAGATTTCCGGCtagagccctcaggcagactgaattctaaatctgtgaaggatcta
aggagacacatgcaccggaaatttccataagccagttgcagttttcaggctaatacagaaaaagatgaacaaacgt
ccagcaagatactttaatcctctattttgctcttccttgtccattgctgctgttaatgtattgctgacctctttcacagttggctct
aaagaatcaaaagaaaaaaactttttatttcttttgctattaaaactactgttcattttgggggctgggggaagtgagcct
gtttggatgatggatgccattccttttgcccagttaaatgttcaccaatcattttaactaaatactcagacttagaagtcaga
tgcttcatgtcacagcatttagtttgttcaacagttgtttcttcagcttcctttgtccagtggaaaaacatgatttactggtctga
caagccaaaaatgttatatctgatattaaatacttaatgctgatttgaagagatagctgaaaccaaggctgaagactgtt
ttactttcagtattttcttttcctcctagtgctatcattagtcacataatgaccttgattttattttaggagcttataaggcatgag
acaatttccatataaatatattaattattgccacatactctaatatagattttggtggataattttgtgggtgtgcattttgttctg
ttttgttgggttttttgttttttttgtttttggcagggtcggtgggggggttatcggaaagaacatgtgagc

Legends:

GCA…GGC: Halo Tag

ccacctgacgtcccaatgat: to overlap pUC57-mini EcoRV cut site for NEB HiFi Assembly atcggaaagaacatgtgagc: to overlap pUC57-mini EcoRV cut site for NEB HiFi Assembly

**Supplementary Material #3: Movie captions**

**Movie S1.**
Movie taken on Zeiss LSM 880 confocal scope showing some purified EGFP-YAP condensates wetting the coverslip (stuck to the bottom of the coverslip).

**Movie S2.**
Zeiss LSM 880 Airyscan movie showing fusion of EGFP-YAP condensates (green) formed after 0.2M sorbitol treatment in HEK293T cell. Movie starts from 1 min after sorbitol treatment.

**Movie S3.**
Zoomed-up movie of nuclear condensates in Movie S2 showing fusion among nuclear EGFP-YAP condensates. Fusion events can be seen from 80 sec to 90 sec, 129 sec to 132 sec, and 150 sec to 156 sec.

**Movie S4.**
Zoomed-up movie of cytoplasmic condensates in Movie S2 showing fusion among cytoplasmic EGFP-YAP condensates. Fusion events can be seen from 0 sec to 8 sec, 16 sec to 21 sec, 73 sec to 100 sec, and 145 sec to 158 sec.

**Movie S5.**
3D ATAC-PALM image of the entire nucleus in control-treated HEK293T cell expressing EGFP-YAP. Accessible chromatin regions labelled by ATAC are mainly diffusive with a few visible small clusters. Imaris software (Bitplane) is used to produce this 3D visualization of the nucleus.

**Movie S6.**
3D ATAC-PALM image of the entire nucleus in 5 min sorbitol-treated HEK293T cell expressing EGFP-YAP. Huge clusters of accessible chromatins labelled by ATAC are seen. Imaris software (Bitplane) is used to produce this 3D visualization of the nucleus.

**Movie S7.**
Colocalization of accessible chromatin clusters labelled by ATAC (red) and EGFP-YAP nuclear condensates (green) in an HEK293T cell at 5 min sorbitol treatment. To visualize accessible chromatin clusters, ATAC-PALM localizations were binned within a cubic of 100 nm with a 3D Gaussian filter and a convolution kernel of $3\times3\times3$. Imaris software (Bitplane) is used to produce this 3D visualization of the two channels.
